# Onset of taste bud cell renewal starts at birth and coincides with a shift in SHH function

**DOI:** 10.1101/2020.10.12.336602

**Authors:** Erin J. Golden, Eric D. Larson, Lauren A. Shechtman, G. Devon Trahan, Dany Gaillard, Timothy J. Fellin, Jennifer K. Scott, Kenneth L. Jones, Linda A. Barlow

## Abstract

Embryonic taste bud primordia are specified as taste placodes on the tongue surface and differentiate into the first taste receptor cells (TRCs) at birth. Throughout adult life, TRCs are continually regenerated from epithelial progenitors. Sonic hedgehog (SHH) signaling regulates TRC development and renewal, repressing taste fate embryonically, but promoting TRC differentiation in adults. Here we show TRC renewal initiates at birth and coincides with onset of SHHs pro-taste function. Using transcriptional profiling to explore molecular regulators of renewal, we identified *Foxa1* and *Foxa2* as potential SHH target genes in lingual progenitors at birth, and show SHH overexpression in vivo alters FOXA1 and FOXA2 expression relevant to taste buds. We further bioinformatically identify genes relevant to cell adhesion and cell locomotion likely regulated by FOXA1;FOXA2, and show expression of these candidates is also altered by forced SHH expression. We present a new model where SHH promotes TRC differentiation by regulating changes in epithelial cell adhesion and migration.

## Introduction

Taste buds are the primary end-organs of the gustatory system, which reside in 3 types of specialized epithelial papillae on the tongue; fungiform (FFP) in the anterior tongue and circumvallate and foliate, posteriorly. Regardless of papilla location, each taste bud houses a collection of heterogeneous taste receptor cells (TRCs) that transduce taste stimuli, including sweet, umami, salt, sour, and bitter, to signal palatability, nutritional value and/or danger of substances in the oral cavity. These signals are conveyed from taste buds to the brain via gustatory nerve fibers of the VII^th^ and IX^th^ cranial nerves. Despite some neuronal characteristics that accompany their sensory function, all TRCs are modified epithelial cells and are continuously renewed (see Barlow and Klein, 2015). Adult taste cells are generated from progenitors adjacent to taste buds that express cytokeratin 14 (KRT14) and KRT5 (Gaillard et al., 2015; Okubo et al., 2009), a population of basal keratinocytes that also gives rise to the non-taste lingual epithelium that covers the tongue surface. Taste-fated daughter cells exit the cell cycle, enter buds as post-mitotic taste precursor cells that express Sonic hedgehog (SHH) and differentiate directly into functional TRCs (Miura et al., 2006; Miura et al., 2014).

Embryonically, taste bud primordia first develop as focal epithelial thickenings, or taste placodes, on the mouse tongue at mid gestation (embryonic day (E) 12.0) (Mistretta, 1972; Mistretta and Liu, 2006). In the anterior tongue, placodes undergo morphogenesis to form FFP, and taste bud primordia are first innervated by gustatory fibers at E14.5 (Lopez and Krimm, 2006); differentiated TRCs are observed in the first postnatal week (Ohtubo et al., 2012; Zhang et al., 2008). Taste placodes express *Shh* at E12.5 (Hall et al., 1999; Jung et al., 1999) and lineage tracing reveals SHH+ placode cells become the first differentiated TRCs after birth (Thirumangalathu et al., 2009). However, SHH+ placode cells do not give rise to adult taste progenitors, as SHH-derived TRCs are steadily lost from taste buds within a few postnatal months (Thirumangalathu et al., 2009). By contrast, lineage tracing of KRT14+ cells initiated in the first postnatal week labels small numbers of taste cells, suggesting at the adult progenitor population is activated in the days following birth (Okubo et al., 2009). It remains to be clarified as to when and how KRT14+ progenitors activate and begin to contribute TRCs to maintain taste buds.

In addition to marking taste placodes in embryonic tongue and post-mitotic precursor cells in mature taste buds, SHH is a key regulator of taste bud development and homeostasis. In embryos, SHH functions to repress taste fate as taste placodes are specified and patterned (El Shahawy et al., 2017; Hall et al., 2003; Iwatsuki et al., 2007; Mistretta et al., 2003); while once taste placodes are established, embryonic taste primordia no longer respond to pharmacological manipulation of Hedgehog signaling (Liu et al., 2004). By contrast, in adults SHH promotes and is required for TRC differentiation. Specifically, ectopic expression of SHH drives formation of ectopic taste buds (Castillo et al., 2014), while pharmacological inhibition or genetic deletion of Hedgehog (Hh) pathway components leads to loss of taste buds (Castillo-Azofeifa et al., 2017; Ermilov et al., 2016; Kumari et al., 2015). Whether the shift in SHH function coincides with progenitor activation has not been explored. SOX2, an SRY-related HMG-box transcription factor, is also a key regulator of embryonic taste bud formation and adult taste cell renewal, and in both contexts, SOX2 is required for TRC differentiation (Castillo-Azofeifa et al., 2018; Okubo et al., 2006). Additionally, SOX2 function is required downstream of SHH, at least in adult taste cell homeostasis.

Here, we sought to define when TRC renewal from adult progenitors occurs, if this renewal coincides with functional shifts in the response of lingual epithelium to SHH, and identify genetic components, in addition to SOX2, that may function downstream of SHH in TRC renewal.

## Results

### Once specified, taste placodes do not receive additional cells from proliferative KRT14+ basal keratinocytes during embryogenesis

Before taste placodes emerge, the tongue anlage is covered by a simple bilayered epithelium that broadly expresses KRT8 (Mbiene and Roberts, 2003). We observed KRT14 co-expression with KRT8 throughout the lingual epithelium at E12.0 (Fig 1A-C’). At E13.5, following placode specification, KRT8 is expressed in placodes and downregulated in non-taste epithelium, although, consistent with previous reports, KRT8 persists in superficial periderm (Fig 1D-F, yellow arrows in E) (Mbiene and Roberts, 2003). At E13.5, KRT14 expression is mostly lost from placodes but evident in cells adjacent to placodes. These patterns are further refined at E16.5; in each FFP, KRT8+ taste bud primordia are KRT14-immunonegative (Fig 1I, asterisk), and KRT8 expression is fully absent from KRT14+ non-taste lingual epithelium as the periderm is lost by this stage (Sengel, 1976) (Fig 1G, H). Thus, taste placodes emerge from embryonic stratifying epithelium that co-expresses KRT14 and KRT8; placode specification results in downregulation of placodal KRT14 and maintenance of KRT8 in taste bud primordia, while KRT14+ non-taste epithelium downregulates KRT8.

**Figure 1.**
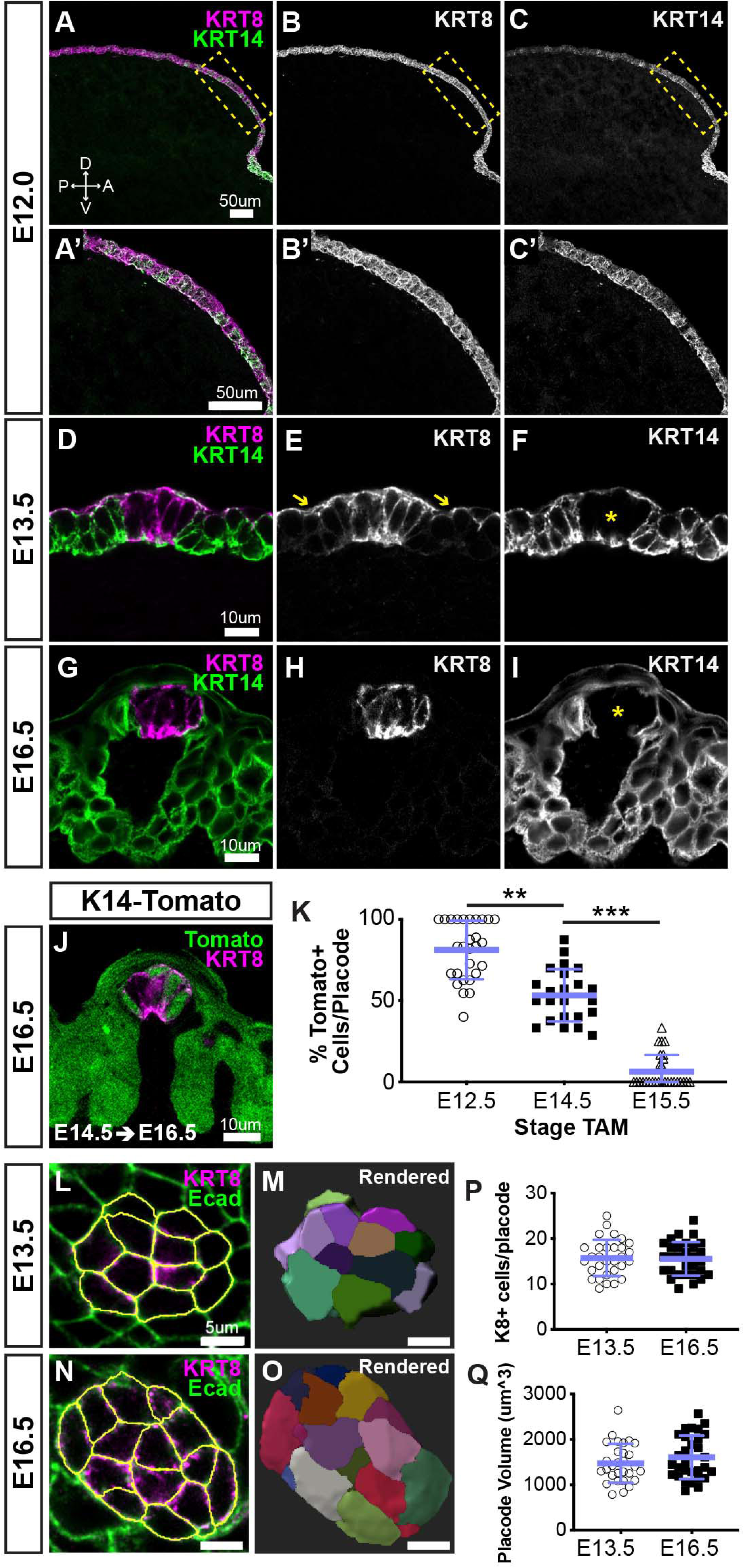
Taste placodes arise from KRT14+ progenitors that do not contribute further to taste bud primordia during embryogenesis. **(A-C’)** Before taste placodes are evident (E12.0), KRT14 (green) and KRT8 (magenta) are co-expressed in lingual epithelium. **(D-F)** At E13.5, KRT8 expression is expressed by taste placodes (asterisk in F), and surface periderm (E, yellow arrows). KRT14 is evident only basally and apically in placodes (asterisk in F) but remains well expressed in non-taste epithelium. (**G-I**) At E16.5, KRT8 is expressed exclusively by taste bud primordia (H), which lack KRT14, while non-taste basal epithelial cells are robustly KRT14+. **(J)** *Krt14*^*CreER*^; *R26R*^*tdTomato*^ (KRT14-Tomato; green) induced at E12.5, labels a subset of KRT8+ (magenta) placode cells at E14.5. (A-J are confocal optical sections acquired at 0.75 µm.**) (K)** Quantification of placodal KRT14-Tomato+ cells 48 hrs after tamoxifen induction at progressive stages, e.g. tamoxifen at E12.5, analysis at E14.5. Blue bars: mean ± SD (n = 3 animals per stage, 6-11 placodes per animal, open and shaded shapes). Student’s *t*-test **p < 0.01, ***p < 0.0001. **(L-O)** E13.5 and E16.5 taste placodes (KRT8+, magenta) in E-cadherin (Ecad) immunostained whole tongues were imaged through their apical-to-basal extent, 3D reconstructed and individual cells defined in Imaris (see methods). White outlines **(L, N)** and randomly assigned colors **(M, O)** indicate individual placode cells. **(P, Q)** Total cell number and placode volume did not differ between stages. Blue bars: mean ± SD (n = 3 animals per stage, 10 placodes per animal, open and shaded shapes).

In adult tongues, TRCs express KRT8 and continuously renew from KRT14+ progenitors, thus we next asked if KRT14+ cells contribute KRT8+ cells to taste placodes and/or taste bud primordia in embryos. Genetic lineage tracing in *Krt14*^*CreER*^;*R26R*^*tdTomato*^ (K14-tomato) embryos was activated via tamoxifen at E12.5, E14.5, or E15.5, spanning the period over which taste placodes are patterned and specified. Embryos were harvested for KRT8 immunostaining after 48-hours. At all stages, tomato+ cells were observed in lingual epithelium (Fig 1J, and data not shown), consistent with KRT14+ keratinocyte generation of lingual epithelium during embryogenesis (Liu et al., 2007). No tomato expression was observed in untreated transgenic embryos (not shown). When K14-tomato was induced at E12.5 and assayed 48 hrs later (E14.5), most KRT8+ placodal cells were tomato+ (Fig 1K). Despite limited KRT14 immunoreactivity in taste placodes at E13.5 (see panel 1F), KRT14-tomato lineage trace initiated at E14.5 resulted in only 50% tomato+/KRT8+ cells 48 hrs later (E16.5. Fig 1J, K). At E17.5 after lineage trace induction at E15.5, most KRT8+ taste bud primordia fully lacked tomato+ cells (Fig 1K).

Because taste placode cells are largely KRT14-immunonegative at E13.5 (Fig 1D-F), we reasoned that tomato-labeling at E16.5 from lineage trace induced at E14.5 (Fig 1J, K) could reflect continued addition of new KRT14+ progenitor-derived cells post-placode specification. Taste placodes are post-mitotic (Farbman and Mbiene, 1991; Mbiene and Roberts, 2003; Thirumangalathu and Barlow, 2015); thus if embryonic KRT14+ progenitors contribute new cells to placodes, the number of cells per placode should increase during development. To determine if cells were added to taste bud primordia after specification, we tallied the cell number and volume of taste placodes and bud primordia at E13.5 and E16.5, respectively (Fig 1L-O and see Methods), and found developing taste buds were static in both cell number and volume (Fig 1P, Q), ruling out an embryonic KRT14 progenitor contribution to taste bud primordia following placode specification. We suspect that low level KRT14 and/or Cre recombinase expression prolonged lineage trace by tamoxifen induction at E14.5.

### KRT14+ progenitors contribute to taste buds at birth

To define the onset of new taste cell production, we first compared the number of KRT8+ cells per taste bud section at E16.5 and early postnatal day (P2, P9) stages. KRT8+ taste cell number was comparable between E16.5 and P2 but increased significantly between P2 and P9 (Fig 2A); suggestive of addition of new taste cells during the first postnatal week. To track entry of cells into buds, we first used thymidine analog birthdating at progressive postnatal timepoints. In pups treated with EdU at P0, EdU+ cells were observed within KRT8+ taste buds 48hr later, indicating addition of new taste cells begins at birth (Fig 2B, C). Comparable rates of intragemmal labeling were observed in postnatal taste buds when EdU was administered at P7 or P14 and quantified at P9 or P16, respectively (Fig 2B, D, E). Previous reports indicated that KRT14 lineage tracing initiated at P2 resulted in sparse labeling of taste bud cells at P9 (Okubo et al., 2009). To confirm new postnatal taste cells were derived from KRT14+ progenitors, we induced Cre recombination in *Krt14*^*CreER*^; *R26R*^*YFP*^ (K14-YFP) pups at P0, P7, or P14 and analyzed YFP+ cell distribution 48 hours later (P2, P9, P16). YFP+ KRT8+ cells were readily observed within taste buds at each stage (Fig 2F-I); additionally, the proportion of taste buds with KRT14-lineage traced cells increased significantly over the first two postnatal weeks (Fig 2F, H, I). Taken together, our results indicate that KRT14+ progenitors generate new taste cells at birth and this contribution steadily increases postnatally, leading to taste bud growth.

**Figure 2.**
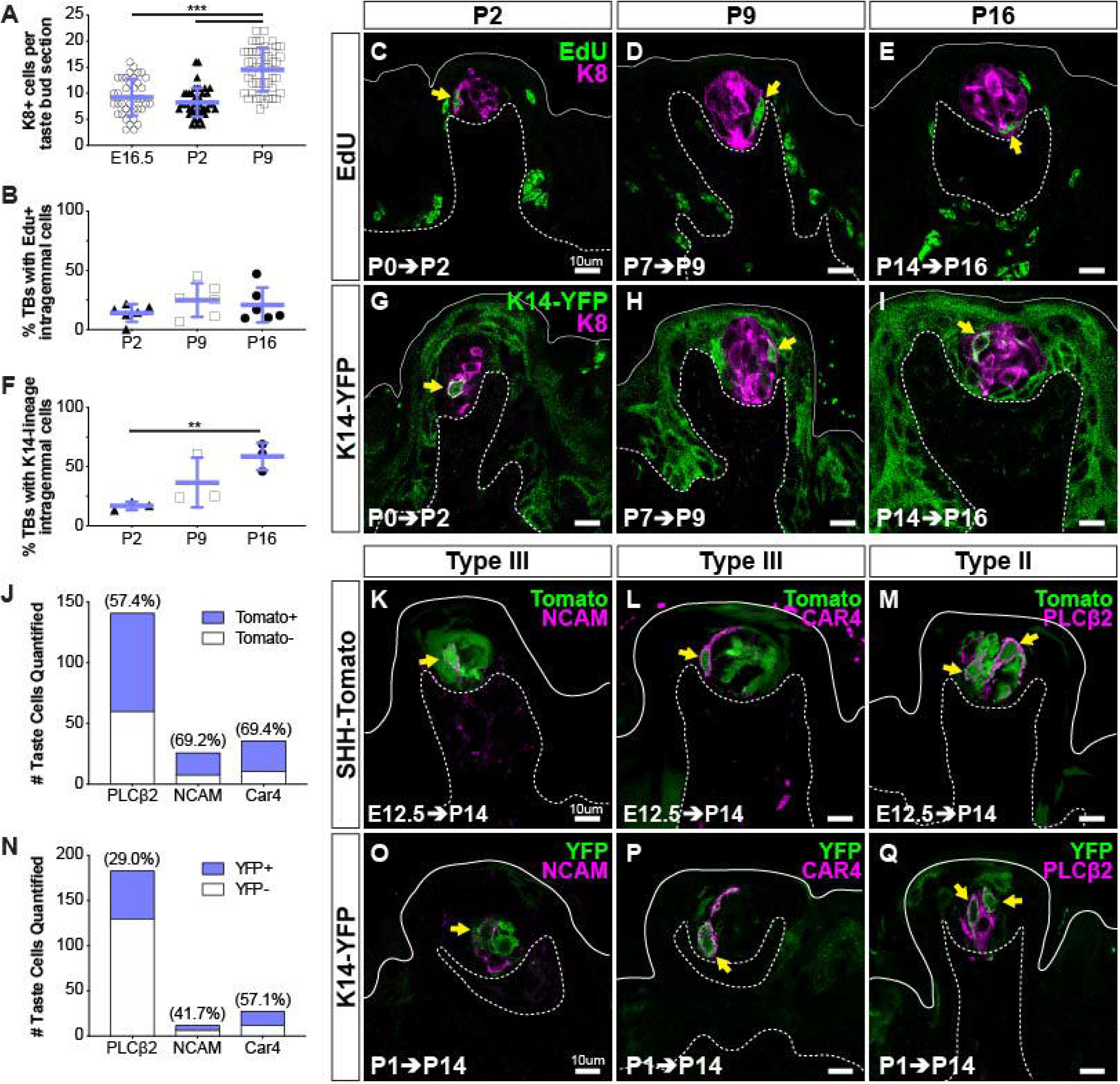
KRT14+ progenitor contribution of new cells to taste buds begins at birth. **(A)** Quantification of KRT8+ cells per FFP section reveals taste bud cell number does not increase until P9. Blue bars: mean ± SD (n = 3 animals per stage, 8-18 taste buds per animal, open and shaded shapes) Student’s t-test ***p<0.001. **(B-E)** In pups that received EdU at P0, P7 or P9, analysis at 48 hrs revealed comparable proportions of taste buds housed newly generated EdU+/KRT8+ cells (yellow arrows in C-E) regardless of postnatal day of labeling (EdU green, KRT8 magenta). (**F-I**) Lineage tracing with *Krt14*^*CreER*^; *R26R*^*YFP*^ (KRT14-YFP) initiated at P0, P7, or P14 assessed at 48 hrs showed extensive YFP expression (green) in FFP non-taste epithelium as well as YFP+/KRT8+ cells in taste buds (magenta, yellow arrows in G-I). **(C-E, G-I)** Dashed lines delimit the basement membrane; solid lines delimit the epithelial surface. **(B, F)** Blue bars: mean ± SD Student’s t-test **p<0.005 (B: n = 6 animals per stage, 14-28 taste buds per animal; F: n = 3 animals per stage, 10-24 taste buds per animal). **(J-M)** SHH+ taste precursor cells are not lineage restricted. *Shh*^*CreER*^; *R26R*^*tdTomato*^ (SHH-Tomato, green) mice treated with TAM at E12.5 reveals similar proportions of type III (NCAM+, CAR4+ magenta in K, L) and type II (PLCß2+ magenta in M) taste cells are tomato+ (green) (N = 3 mice, counts from 24 NCAM+, 30 CAR4+ and 30 PLCß2+ total TBs). (**N-Q**) Postnatally activated KRT14+ progenitors are not lineage restricted. *Krt14*^*CreER*^;*R26R*^*YFP*^ (K14-YFP, green) mice treated with TAM at P1 labels both type III (NCAM+, CAR4+ magenta in O, P) and type II (PLCß2 magenta in Q). Double labeled cells indicated with yellow arrows in all image panels. Dashed lines delimit the basement membrane; solid lines delimit the epithelial surface.

### Embryonic Shh+ taste placodes and postnatal KRT14+ progenitors give rise to comparable proportions of differentiated taste cell types

Murine taste buds each house ∼60 functionally and morphologically heterogeneous cells, including type I support cells, type II detectors of sweet, bitter or umami, and type III sour receptor cells (Roper and Chaudhari, 2017). In adult taste buds, SHH+ cells are immediate precursors for all mature TRC types (Miura et al., 2014). Whether SHH+ embryonic taste precursors are similarly competent has not been fully defined. Previously, we used a low efficiency Cre reporter to trace SHH-expressing taste placode cells from E12.5 into adulthood (*Shh*^*CreERT2*^; *R26R*^*LacZ*^) (Thirumangalathu et al., 2009). At 6 weeks, embryonically derived type I and type II TRCs were readily observed, but lineage labeled type III cells were not, suggesting embryonic taste precursors may be lineage restricted. However, type III cells account for <10% of adult TRCs (Ma et al., 2007; Ohtubo and Yoshii, 2011) and our previous experimental parameters were not optimized to detect this less common cell population.

Here, to determine if type III, like type I and II TRCs, arise from Shh+ placodes, we employed a high efficency Cre reporter allele, *R26R*^*tdTomato*^ (Madisen et al., 2010) to assess taste cell fate in *Shh*^*CreERT2*^; *R26R*^*tdTomato*^ (Shh-tomato) pups. Cre induction at E12.5 resulted in robust lineage labeling at P14: 95.2% (± 0.01% s.e.m., N = 3 animals) of taste buds were tomato+ with 10.6 (± 1.1 s.e.m.) tomato+ cells per taste bud profile (compare with our previous finding of 49.7% ± 7.83% of labeled taste buds with 2.9 ± 0.41 placode-descendent cells per TB profile) (Thirumangalathu et al., 2009).

Immunostaining for type III TRC markers NCAM and CAR4 revealed significant double labeling in Shh-tomato taste buds. Specifically, ∼70% of both NCAM+ cells and CAR4+ cells were tomato+ at P14 (Fig 2J-L). Similarly, ∼60% of PLCß2+ type II cells were tomato+ at P14, following placodal lineage trace initiated at E12.5 (Fig 2J, M). Together with our previously published findings (Thirumangalathu et al., 2009), we show Shh+ taste placode cells give rise to all TRC types postnatally. Further, we find placodally derived cells comprise slight majorities of type II and III TRCs at P14, suggesting unlabeled TRCs were new cells derived from KRT14 progenitors in the first postnatal weeks.

To test this, we induced K14-YFP lineage trace at P0, P7, or P14 and following a 48hr or 72hr chase, immunostained tongue sections for markers of type II and III TRCs as above. However, despite ample numbers of YFP+ cells within taste buds, we did not detect YFP+ TRCs immunopositive for PLCß2, NCAM or CAR4 (data not shown). In adult rodents, type II and III TRCs require ∼3.5 days from their terminal division to turn on expression of cell type-specific immunomarkers (Cho et al., 1998; Hamamichi et al., 2006; Perea-Martinez et al., 2013); the lack of double labeling in short term postnatal lineage tracing here suggested the rate of differentiation of postnatally derived TRCs is comparable to that of adults. Thus, we next examined the fate of newly generated TRCs in tongues from mice traced by induction of K14-YFP at P1 and harvested at P15. Similar to *Shh*^*CreER*^ lineage tracing, KRT14 lineage tracing resulted in labeling of both type II and III cells in comparable but not identical proportions (Fig 2N-Q), indicating that SHH+ taste primordia and postnatal KRT14+ progenitors, nonetheless, have no bias in terms of taste cell type production and are both competent to differentiate the full taste bud lineage.

### Shh represses taste fate during placode specification and promotes taste fate postnatally

The Hh pathway has opposing roles in embryonic versus adult taste epithelium – restricting taste fate during placode specification, and promoting taste cell differentiation during adult renewal (Barlow, 2015). We next used genetic deletion and activation of the Shh pathway to determine when this shift occurs.

Pharmacologic inhibition or genetic deletion of Shh signaling during placode specification results in overproduction of enlarged taste buds (El Shahawy et al., 2017; Hall et al., 2003; Iwatsuki et al., 2007; Liu et al., 2004; Mistretta et al., 2003). However, the effects of this disruption on later aspects of taste bud development, including TRC differentiation, have not been assessed. Previously, we observed differentiated Type I and II TRCs within a subset of taste buds at E18.5 (Thirumangalathu and Barlow, 2015) and further analysis here reveals differentiation of all three TRC types is underway at E17.5 (Fig S1). To determine if loss of Shh signaling affects TRC differentation, we induced *Shh* deletion in *Shh*^*CreERT2/fl*^ (Shh-ShhcKO) mice (Castillo-Azofeifa et al., 2017; El Shahawy et al., 2017). Tamoxifen induction at E13.5 resulted in reduced *Shh* mRNA in E18.5 lingual epithelium, as well as reduced expression of its target transcriptional regulator, *Gli1* (Fig S2A). In Shh-ShhcKO embyos induced at E12.5 and analyzed E18.5, the total number and size of KRT8+ taste buds and FFP were significantly increased (Fig S2B-F). Additionally, E12.5 Shh-ShhcKO significantly increased the proportion of taste buds containing differentiated type I (NTPdase2+), II (PLCß2+), and III (NCAM+) TRCs compared to *Shh*^*wt/fl*^ littermate controls (Fig S2G-I), indicating that Shh reduction during placode specification accelerated taste bud differentiaton. This outcome is likely due to the loss of Shh’s repressive role on Wnt/ß-catenin signaling during placode specification that in turn leads to precocious TRC differentiation (Iwatsuki et al., 2007; Thirumangalathu and Barlow, 2015)

Pharmacologic inhibition of the Shh pathway effector, Smoothened, in embryonic rat tongue explants suggests Shh no longer regulates placode development once these structures have formed (Liu et al., 2004). Consistent with this, ShhcKO induced at E16.5 did not alter the number or size of FFP or KRT8+ taste buds (Fig S2B, J, K). Moreover, E16.5 induction of ShhcKO did not acclerate TRC differentiation; similar proportions of taste buds housed type I (NTPdase2+), II (PLCß2+), and III (NCAM+) TRCs in ShhcKO and control tongues at E20 (Fig S2L-N). Our in vivo results confirm and expand upon in vitro studies showing taste epithelium is not sensitive to Shh pathway inhibition in late gestation (Liu et al., 2004) (but see Cohen et al., 2019).

In contrast to its repressive function in embryos, in adult lingual epithelium misexpression of SHH induces ectopic taste buds in non-taste lingual epithelium (Castillo et al., 2014). Thus, we next asked if Shh promotes taste fate at birth, coinciding with the onset of taste cell production by KRT14+ progenitors. *Krt14*^*CreER*^;*Rosa*^*SHHcKI-IRES-nVenus*^ (K14-SHHcKI) was induced in neonatal mice and tongues analyzed after 14 days. In addition to SHH, *Rosa*^*SHHcKI-IRES-nVenus*^ drives expression of nuclear Venus (YFP) in SHH+ cells. As in adults (Castillo et al., 2014), neonatal *Krt14*^*CreER*^ recombination resulted in patches of SHH-YFP+ cells in both FFP and non-taste epithelium. When K14-ShhcKI was induced at P1 and assessed at P15, small numbers of ectopic KRT8+ cells were detected in SHH-YFP+ non-taste lingual epithelium (Fig 3A, B). P1 induction of K14-ShhcKI largely resulted in individual ectopic KRT8*+* cells (Fig 3A, yellow arrow). Significantly more and larger KRT8+ cell clusters formed in K14-ShhcKI tongues induced at P14 and harvested at P28 (Fig 3C, D), and these ectopic buds co-expressed markers of type I (NTPdase2), II (PLCß2), and III TRCs (5HT; (Huang et al., 2005)) (Fig 3E-G). Notably, forced SHH expression postnatally had no effect on endogenous FFP taste bud number (Fig. 3B, D). By contrast, embryonic induction of ShhcKI at E12.5 did not lead to formation of ectopic buds at P14, but as expected repressed endogenous FFP taste bud development (Fig S3).

**Figure 3.**
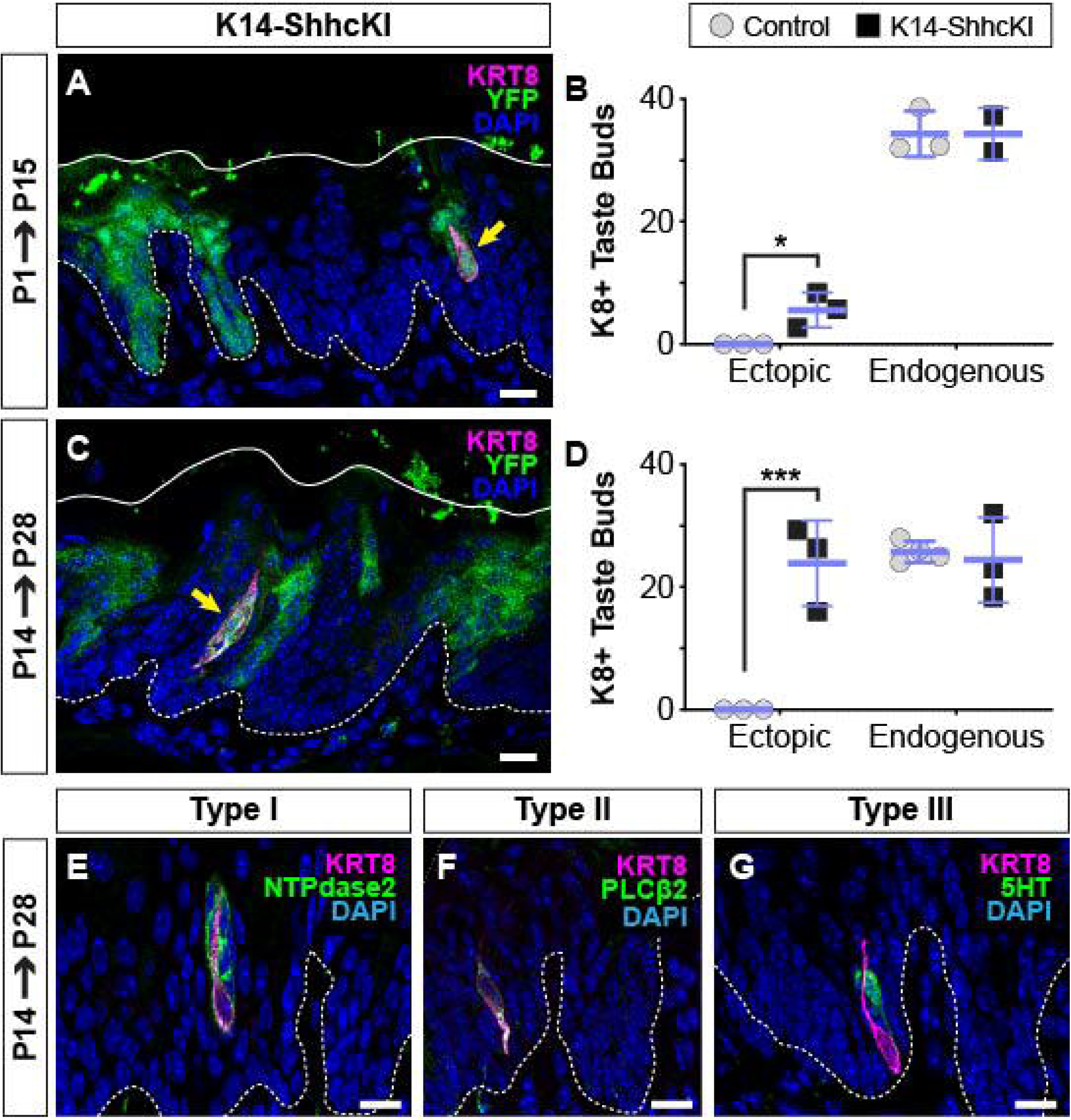
SHH promotes taste fate at birth. **(A, B)** KRT14-SHHcKI-YFP induction at P1 drives production of small numbers of ectopic KRT8+ cells (magenta, yellow arrow) in patches of YFP+ non-taste lingual epithelium at P14 but does not impact the number of endogenous taste buds (n = 3 mice per timepoint) Student’s t-test *p<0.03. **(C, D)** KRT14-SHHcKI-YFP induction at P14 induces more and larger ectopic KRT8+ taste cell clusters at P28, with endogenous taste bud number unaffected (n = 3 mice per timepoint) ***p<0.005 Student’s t-test. (**E-G**) Ectopic taste buds at P28 house cells expressing markers of type I (NTPdase2 **E)**, II (PLCß2, **F)**, and III (5HT, **G)** taste cells. (**A, C, E-G**) Dashed lines delimit the basement membrane; solid lines delimit the epithelial surface; yellow arrows indicate ectopic KRT8+ taste cells. Nuclei counter stained with DAPI. Scale bar: 10 µm

### Neurally supplied SHH is not required for taste progenitor activation

In adult tongue, SHH is expressed by postmitotic taste precursor cells and gustatory sensory neurons of the VII^th^ cranial ganglion, which together support continual TRC differentiation (Castillo-Azofeifa et al., 2017; Lu et al., 2018; Miura et al., 2003). However, embryonic taste placodes are not innervated at specification, hence the only early source of lingual SHH is placodal (Hall et al., 1999). Gustatory neurons develop by E10.5 (Cordes, 2001) and gustatory neurites first penetrate the taste epithelium at E14.5 (Krimm, 2007; Lopez and Krimm, 2006). When gustatory neurons start to express SHH has not been determined. Thus, we next asked whether SHHs functional shift from a repressor of taste fate to a driver of TRC differentiation is correlated with the onset of gustatory neuron SHH expression.

Lineage tracing with *Shh*^*CreER*^;*R26T*^*tdTomato*^ leads to ample labeling of gustatory innervation in adult mice after 2 weeks (Castillo-Azofeifa et al., 2017). Thus, we induced Shh-tomato lineage tracing in embryos during placode specification (E12.5) and quantified the number of tomato+ nerve fibers within the core of each FFP at P14. In addition to ample taste bud labeling as expected, Shh lineage tracing revealed ∼1/3^rd^ of taste buds were contacted by one or more tomato+ neurites at P14 (Fig 4A, C, D). Thus, some gustatory neurons already express Shh as early as E12.5. When lineage tracing was induced at birth (P0), well after nerve contacts are established, and examined at P14, the proportion of FFP with tomato+ neurites was significantly increased and taste buds were contacted by more tomato+ neurites (Fig 4B-D). Taken together, these results indicate that some gustatory neurons express SHH embryonically, well before the shift in SHH function, which occurs at birth.

**Figure 4.**
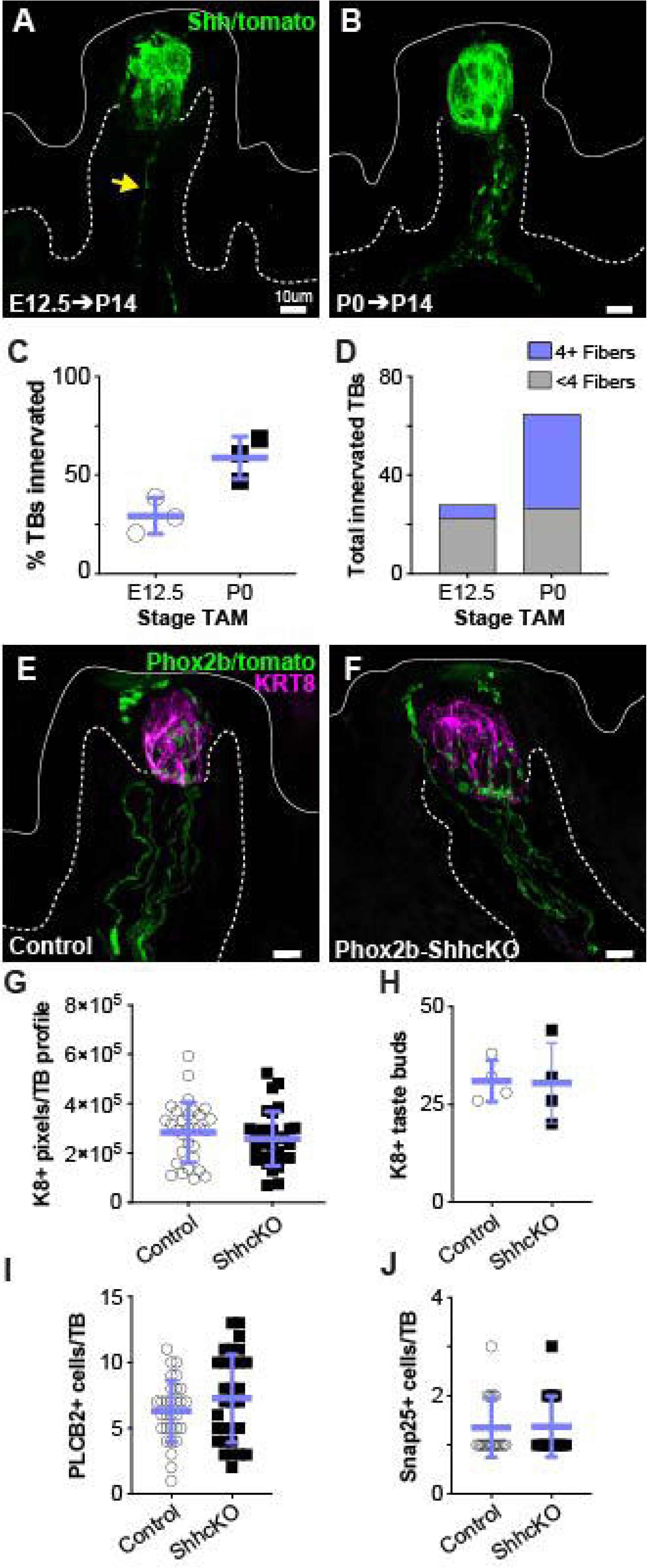
Embryonic gustatory neurons express SHH, but neural SHH is not required for taste development or postnatal taste bud maintenance. **(A)** *Shh*^*CreER*^; *R26R*^*tdTomato*^ (SHH-tomato) induction at E12.5 drives tomato expression (green) in taste cells and sparse gustatory neurites (yellow arrow) at P14. **(B)** Induction of Shh-tomato at P0 results in tomato+ taste cells and more numerous reporter-expressing gustatory neurites at P14. **(C)** More taste buds are innervated at P14 by P0 SHH-tomato lineage traced fibers compared to fibers lineage traced starting at E12.5. Student’s *t-*test p=0.02. **(D)** At P14, individual taste buds are innervated more densely when lineage trace is initiated at P0 compared to initiation at E12.5 (n = 3 mice per stage, E12.5 induction - 98 taste buds, P0 induction – 111 taste buds). **(E, F)** *Phox2b*^*Cre*^ drives tomato expression (green) in gustatory neurites innervating KRT8+ taste buds (magenta) in control (*Phox2bCre;R26R*^*tdTomato*^) and Phox2b-SHHcKO (*Phox2bCre;Shh*^*fl/fl*^;*R26R*^*tdTomato*^) mice. **(G, H)** Taste bud size (KRT8+ pixels per taste bud profile) and number do not differ between controls and Phox2b-SHHcKO mice **(I-K)** Phox2b-SHHcKO does not disrupt differentiation of PLCß2+ type II **(I)** or SNAP25+ type III **(J)** taste cells. Blue bars: mean ± SD (n = 3-4 mice per condition; 10 taste buds per animal; open and shaded shapes).

We next tested if neuronal SHH is required for embryonic taste bud development and/or progenitor function postnatally. PHOX2b is a transcription factor expressed by developing cranial sensory neurons, including gustatory neurons that innervate FFP taste buds (Ohman-Gault et al., 2017; Sajgo et al., 2016). We used *Phox2b*^*Cre*^ (Scott et al., 2011) to constitutively delete SHH and drive tomato expression within PHOX2b+ neurons (*Phox2b*^*Cre*^;*Shh*^*fl/fl*^;*R26R*^*tdTomato*^) (Phox2b-ShhcKO) (Fig 4E, F). When assessed at 10 weeks, taste buds in Phox2b-ShhcKO mice were indistinguishable from control *Phox2b*^*Cre*^; *R26R*^*tdTomato*^ littermates; we found no differences in taste bud number, size (KRT8+ pixels/taste bud), or type II or III TRC differentiation (Fig 4G-J). These results indicate that neural SHH is dispensable for embryonic taste placode specification and postnatal taste bud differentiation, and are consistent with previously published findings in adults where epithelial SHH can compensate for loss of neuronal SHH to maintain adult taste buds (Castillo-Azofeifa et al., 2017).

### Transcriptional profiling of lingual progenitors reveals additional candidates that function downstream of SHH in TRC renewal

The transcription factor SOX2 plays a critical role in taste bud development and renewal, and is required downstream of SHH for adult taste bud differentiation (Castillo-Azofeifa et al., 2018; Okubo et al., 2006). At P0, SOX2 is highly expressed in and around KRT8+ taste buds (Fig 5A, yellow arrows), moderately expressed by FFP basal progenitors (Fig 5A yellow arrowheads), and dimly expressed by basal keratinocytes of the interpapilliary non-taste epithelium (Fig 5A, white arrowheads) (and see (Nakayama et al., 2015)); an expression pattern that is recapitulated in *SOX2-EGFP* (SOX2-GFP) reporter mice (Fig 5B) (Okubo et al., 2006; Taranova et al., 2006). This expression pattern has led to the suggestion that high SOX2 is associated with taste fate, while moderate/dim SOX2 marks non-taste progenitors (Castillo-Azofeifa et al., 2018; Nakayama et al., 2015; Okubo et al., 2006). Thus, we sought to compare the transcriptional profiles of SOX2-GFP high vs low expressing epithelial cell populations at P0 to identify genetic pathways associated with activation of KRT14+ progenitors.

**Figure 5.**
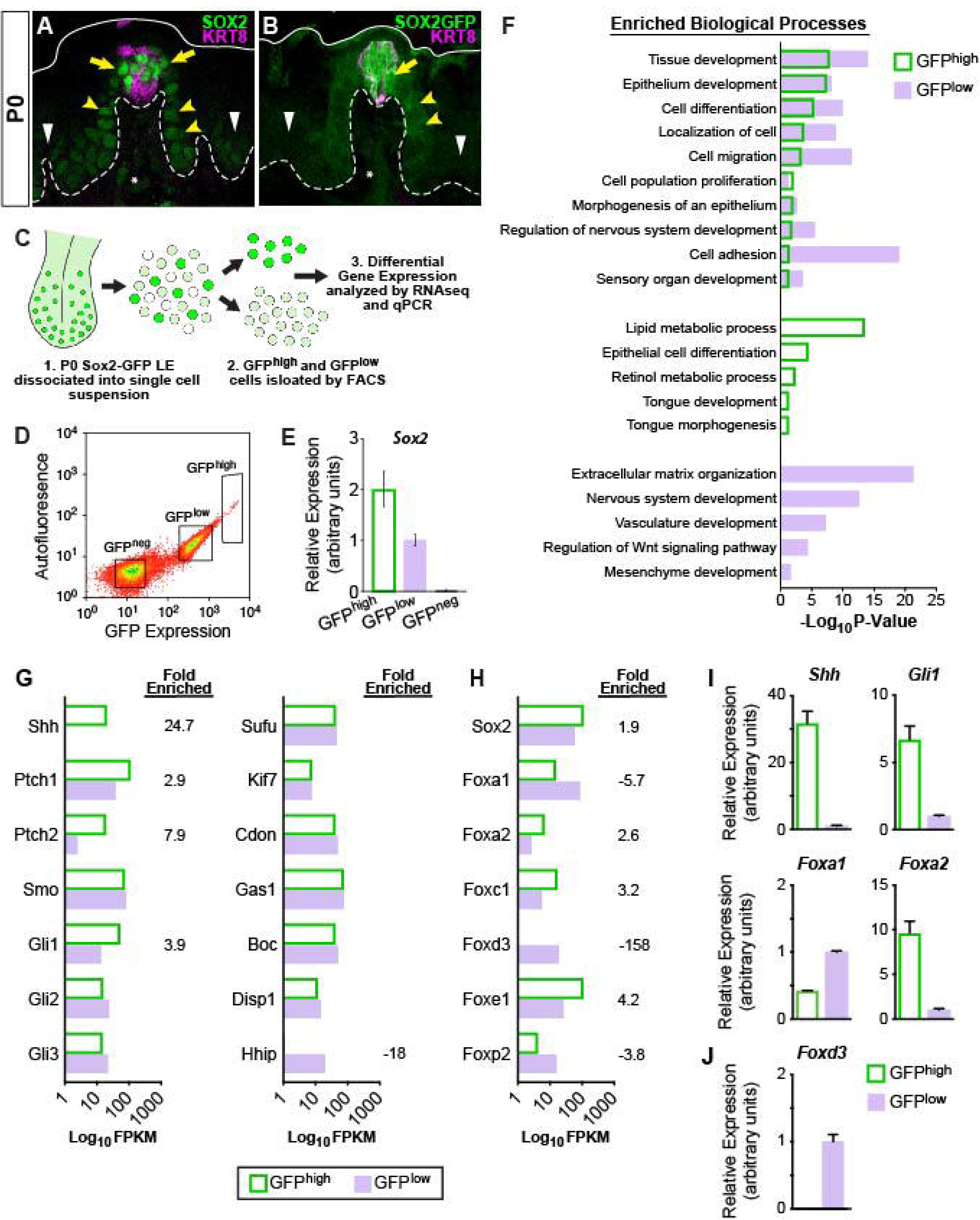
Shh pathway associated genes are differentially expressed in SOX2-GFP lingual epithelial cells. **(A, B)** At P0, SOX2 and SOX2-GFP (green) are expressed highly in KRT8+ taste buds (magenta) and perigemmal cells (yellow arrows), at lower levels in FFP epithelium (yellow arrowheads) and least intensely in non-taste epithelial basal cells (white arrowheads). Dashed lines delimit basement membrane, solid lines delimit apical epithelial surface. (**C**) Experimental procedure to isolate SOX2-GFP^high^ and SOX2-GFP^low^ lingual epithelial cells of P0 pups for RNAseq and qPCR analysis. (**D**) SOX2-GFP^high^, SOX2-GFP^low^ and SOX2-GFP^neg^ FAC sorted lingual epithelial cells were collected in discrete fluorescence bins from an expression continuum. (**E**) qPCR for *Sox2* confirms expression is highest in SOX2-GFP^high^ cells and absent in SOX2-GFP^neg^ cells. (**F**) GO term analysis of differentially expressed genes revealed processes enriched in SOX2-GFP^high^ vs SOX2-GFP^low^ populations (see text for details). (**G**) SHH pathway genes are enriched in SOX2-GFP epithelial cells. (H) Transcription factors associated with SHH signaling are differentially expressed in SOX2-GFP^high^ v SOX2GFP^low^ epithelial cells. (**I**) qPCR for SHH pathway associated genes confirms *Shh, Gli1*, and *Foxa2* are upregulated in SOX2-GFP^high^ cells, while *Foxa1* is more highly expressed in SOX2-GFP^low^ cells. (**J**) *Foxd3* is expressed only in the SOX2-GFPlow cells, consistent with the inclusion of mesenchymal cells in this population.

Dissociated SOX2-GFP lingual epithelial cells from 65 pups (10 litters) were FAC sorted into GFP^high^ and GFP^low^ populations (Fig 5C, D), and RNA isolated from ∼100,000 cells per sorted cell population for gene profiling. Confirming the efficacy of cell sorting, *Sox2* was expressed 2x higher in GFP^high^ cells compared to GFP^low^ cells by qPCR (Fig 5E). To identify candidate genes regulating epithelial cell fate we concentrated our RNAseq analyses on differentially expressed protein-coding genes (DEGs) enriched in GFP^high^ or GFP^low^ cells (Supplementary Tables S1-S3). Using a 5 FPKM threshold and enrichment of 1.5-fold or greater expression over the opposing population, we identified 1032 GFP^high^ and 970 GFP^low^ DEGs (Fig S4, Tables S2, S3). As expected, taste and progenitor-associated genes were enriched in GFP^high^ cells, including *Slc6a11, Shh, Prox1, Lrmp, Ptch2, Slc6a13, Lgr5, Rarb, Six1, Bdnf, Wnt10b, Krt8, Krt19, Krt7, and Hes6* (Table S2). The top 50 GFP^high^ DEGs also included genes broadly associated with epithelial development; *Gpa33, Stfa1, Krt75, Krt4*, and *Elf5* (Fig S3). By contrast GFP^low^ DEGs included markers of both epithelium (*Lyg1, Krtap3-1, Krt84, Krt36, Krt83, Krtap13-1, Gfra2, Itga8, Krt81*) and connective tissue (*Foxd3, Sox10, Mpz, Ednrb, Zeb2, Cdh6*) (Fig S4, Table S3). As some lingual mesenchymal cells appear to express low levels of SOX2 or Sox2-GFP (Fig 5A, B asterisks, and see (Arnold et al., 2011), these cells were likely included in the SOX2-GFP^low^ population. Thus, GFP^high^ cells comprise an epithelial taste/progenitor enriched population, while GFP^low^ cells represent a mixed population of lingual epithelium and connective tissue. These conclusions were further supported by Gene Ontology (GO) analyses (Fig 5F, Tables S4, S5). GO terms for both populations included biological processes associated with developing epithelia (i.e. *Epithelium development, Cell migration, Cell adhesion*). GFP^low^ analyses also included terms for non-epithelial tissues *(Vasculature development, Mesenchyme development*). While both datasets triggered the term *Sensory organ development*, only the GFP^high^ dataset was associated with the system-specific terms *Tongue development* and *Tongue morphogenesis*. In all, SOX2-GFP-based FACS allowed for efficient isolation of differentiating taste and progenitor cells for gene expression profiling.

As expected, SHH pathway components were expressed by both GFP^high^ and GFP^low^ cells (Fig 5G). Consistent with previous reports, *Shh, Ptch1*, and *Gli1* were enriched in GFP^high^ compared to GFP^low^ cells (Fig 5G, I) (Hall et al., 1999; Liu et al., 2013). Additionally, the transmembrane receptor *Ptch2* was highly enriched in the GFP^high^ population, while Hedgehog-interacting protein *Hhip* was highly enriched in GFP^low^ cells. The role of these latter hedgehog pathway genes has not been explored in taste epithelium.

To identify specific candidate transcriptional regulators acting downstream of SHH within P0 lingual epithelium, we performed unbiased motif enrichment analysis on the GFP^high^ DEG dataset. Regions of interest (ROI), which included 1000kb upstream and 200bp downstream of the transcription start site (TSS), were generated for each gene and common regulatory sequences were evaluated in MEME-Suite AME (McLeay and Bailey, 2010). Of the 201 transcription factors identified by AME, only 123 were expressed within our P0 dataset (Table S6). A literature search revealed ∼20% of these candidates were associated directly or indirectly with the SHH pathway; including Sox2, FoxA1, and FoxA2.

In our dataset *Sox2* and *Foxa2* were enriched within the GFP^high^ population (1.9x and 2.6x, respectively), while *Foxa1* was enriched in GFP^low^ cells (−5.65x) (Fig 5H, I). Notably, *Foxd3*, a canonical mesenchyme gene, was expressed exclusively in GFP^low^ cells (Fig 5H, J). From published reports, FOXA2 and FOXA1 are broadly expressed in developing lingual epithelium at E13.5 - 15.5, and FOXA1 maintains broad expression at E18.5; by contrast, at E18.5 FOXA2 expression is absent from the general tongue epithelium and restricted to subsets of cells within taste buds and to clusters of keratinocytes at the base of FFP (Besnard et al., 2004; Luo et al., 2009). To more precisely define the spatial expression of FOXA1 and FOXA2 in developing taste vs non-taste epithelium, we compared SOX2, FOXA2 and FOXA1 expression at E13.5 during placode specification when SHH signaling represses taste fate, E16.5 when taste development is SHH-insensitive, and P0 as taste cell renewal begins and SHH promotes taste fate.

At E13.5, SOX2 is more highly expressed in KRT8+ taste placodes than by surrounding non-taste epithelium (Nakayama et al., 2015; Okubo et al., 2006) (Fig S5A, asterisk and arrowheads). FOXA2 and FOXA1 are both expressed broadly by taste placode and non-taste epithelium (Fig. S5B, C. placode: asterisk, non-taste epithelium: arrowheads). At E16.5, SOX2 expression remains robust in taste primordia and perigemmal cells adjacent to buds (Fig S5D, yellow arrows) but is lower in FFP and non-taste epithelial basal cells (white arrows and arrowheads). At E16.5 FOXA2 expression has altered, and is most intense in KRT8+ taste cells, moderate at the base of each FFP, and dim in non-taste epithelium (Fig S5E white arrows and arrowheads); FOXA2 expression is absent from perigemmal cells (Fig S5E yellow arrowheads). FOXA1 expression at E16.5 has also shifted and is evident at low levels in sparse taste cells, absent in perigemmal cells (Fig S5F yellow arrowheads), but robust in the developing filiform papilla and interpapillary non-taste epithelium (fi, ip). At P0, the pattern of SOX2 and FOXA2 in taste buds and FFP was mostly unchanged from E16.5 (Fig S5G, H), although FOXA2 was significantly diminished in non-taste epithelium. FOXA1 expression remained low in subsets of taste cells and largely lacking in FFP keratinocytes, but strong in filiform papillae and interpapillary epithelium (Fig S5I). We conjecture that changes in expression of these 3 transcription factors could contribute to the transformation of the lingual epithelial response to SHH from an embryonic repressor to a taste fate promoter in adults.

Induction of K14-ShhcKI in adult lingual epithelium leads to formation of ectopic taste buds, including induction of high SOX2 within and moderate SOX2 expression around ectopic buds (Castillo et al., 2014). To determine if changes in FOXA1 and FOXA2 expression also occur, we induced K14-ShhcKI at P1 and analyzed *Sox2, Foxa2* and *Foxa1* expression at P15. Quantitative PCR of *Shh* and *Gli1* revealed hedgehog signaling was highly elevated in K14-ShhcKI tongues compared to controls, as was *Sox2* (Fig 6A). Likewise, *Foxa2* was highly upregulated by K14-ShhcKI. However, *Foxa1* was significantly reduced by SHH overexpression (Fig 6B).

**Figure 6.**
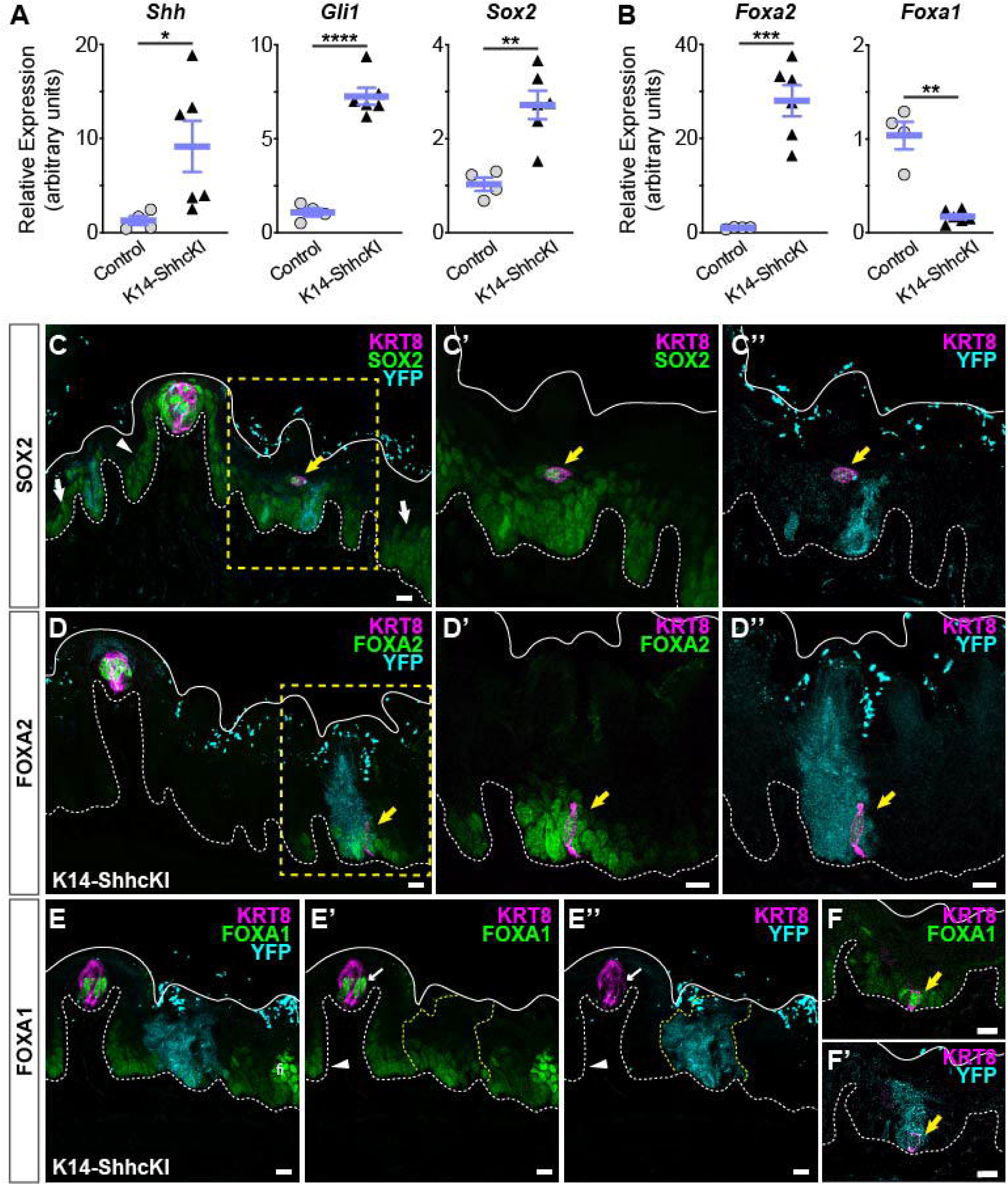
SOX2, FOXA2, and FOXA1 expression in lingual epithelium are altered by postnatal induction of SHH. **(A**) *Shh, Gli1* and *Sox2* are increased in lingual epithelium harvested from P14 *Krt14*^*CreER*^;*ShhcKI-YFP* (KRT14-SHHcKI) pups induced with tamoxifen at P0. (**B**) *Foxa2* expression is increased in SHHcKI epithelium while *Foxa1* is significantly reduced. A, B: n = 4 - 6 mice per genotype, Student’s t-test *p < 0.05; **p < 0.01; *** p < 0.001; **** p < 0.0001. (**C-C”**) SOX2 expression (green) is upregulated in and around patches of SHH-YFP+ cells (teal) in lingual epithelium of P14 mice induced with tamoxifen at P0. Ectopic KRT8+ taste cells (magenta) express elevated SOX2 (yellow arrow). (**D-D”**) FOXA2 is upregulated in and around patches of SHHcKI-YFP+ cells and is detected in ectopic KRT8+ taste cells (yellow arrow). (**E-E”**) FOXA1 expression appears reduced in and around patches of SHHcKI-YFP+ cells (dashed yellow lines) but is also detected in occasional ectopic KRT8+ taste cells in SHHcKI-YFP+ domains (**F, F’**). Scale bars: 10µm. Basement membrane delimited by dashed white line; solid white lines mark epithelial surface.

As expected K14-ShhcKI induction at P0 resulted in mosaic expression of SHH-YFP+ cell patches throughout an otherwise normal lingual epithelium at P15. In control regions of mutant tongues, SOX2 expression at P15 resembles that reported in adult mice (Fig 6C; FFP basal keratinocytes, arrowhead), and non-taste basal keratinocytes (white arrows) (Castillo-Azofeifa et al., 2018; Okubo et al., 2006; Suzuki, 2008). In and around SHHcKI-YFP domains, SOX2 expression was detected in ectopic KRT8+ cells (yellow arrow) surrounded by dim SOX2+ cells (Fig 6C’, C”), similar to the epithelial response observed in adult mice (Castillo et al., 2014).

At P15 in control domains, FOXA2 is strongly expressed by KRT8+ taste cells (Fig 6D). In response to SHH induction, like SOX2, FOXA2 expression was increased in and around patches of ectopic SHH-YFP+ cells, including signal in ectopic KRT8+ taste cells (Fig 6D-D”, yellow arrow). In control epithelium, FOXA1 is expressed by small numbers of taste cells, and keratinocytes of filiform papillae and interpapillary epithelium (Fig 6E-E’). FOXA1 expression appeared diminished in and around SHH-YFP+ patches (Fig. 6E’, E” yellow dash), consistent with our qPCR results (see Fig 6B), but was observed in occasional ectopic KRT8+ taste cells (Fig 6F, F’).

Given expression of these 3 transcription factors is affected by SHH, we next used motif analysis to identify potential target genes that may function downstream of SOX2, FOXA2 and/or FOXA1 in TRC differentiation. Within the SOX2-GFP^high^ dataset, 45.6% of DEG (differentially expressed gene) ROIs (regions of interest) contained a SOX2 binding motif, 38.4% a FOXA2 motif and 24.8% a FOXA1 motif; 143 DEG ROIs contained motifs of all three transcription factors while 90 DEGs had motifs for SOX2 and FOXA2, 102 DEGs had motifs for FOXA1 and FOXA2, and only 6 DEGs had motifs for SOX2 and FOXA1 (Fig 7A Table S7). We next asked what biological processes might be regulated by these different combinations of transcription factors using gene ontology analysis of the short list of DEGs with motifs for each combination of SOX2, FOXA2 and/or FOXA1 (Table S8). No significant biological GO terms were associated with genes regulated by only FOXA2 or FOXA1, the combination of SOX2;FOXA1, or all 3 genes. Both SOX2 and SOX2;FOXA2 DEGs were mildly enriched for a small number of developmental and metabolic processes. However, FOXA2;FOXA1 DEGs were strongly enriched for terms including: *Regulation of cell adhesion* and *Positive regulation of cell migration* (Table S9). Among genes common to both GO terms were *Pdpn* (*Podoplanin*), *Plet1* (*Placenta expressed transcript 1*), *Cxcl10* and *Tgfb2*; while *Runx1, Fbln2* (*Fibulin 2*) and *Ephb4* were unique to *Cell adhesion*. Our *in silico* results suggested these genes may function in taste progenitor production of new taste cells at P0.

**Figure 7.**
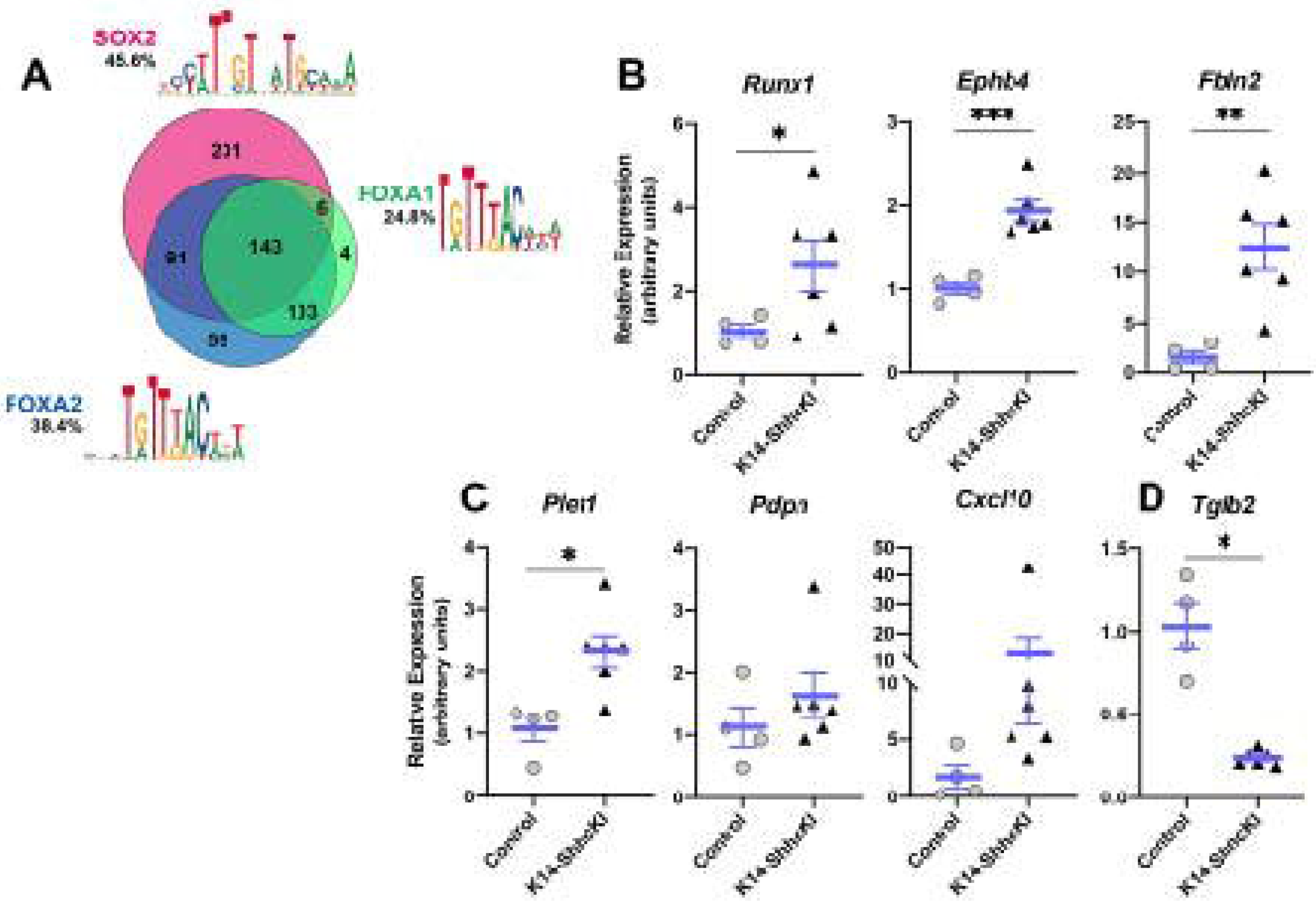
Expression of FOXA2;FOXA1 putative target genes associated with cell adhesion and movement is altered in response to forced SHH expression. (**A**) Motif analysis of DEGs in SOX2-GFP+ cells identified candidate target genes of SOX2, FOXA2 and FOXA1 in lingual epithelium at P0. See text for details **(B-D)** Expression of putative target genes of FOXA2 and FOXA1 is altered in P14 lingual epithelium from KRT14-SHHcKI mice induced at P0. (**B**) *Runx1, Ephb4* and *Fbln2* are significantly upregulated in KRT14-SHHcKI epithelium compared to controls. (**C**) *Plet1* is upregulated and *Pdpn* and *Cxcl10* trend upward following KRT14-SHHcKI induction. (**D**) *Tgfb2* is significantly downregulated by KRT14-SHHcKI. B-D: n = 4 - 6 mice per genotype, Student’s t test, * p<0.05, ** p<0.005, *** p<0.0005, ****.

We next assessed if expression levels of these genes were altered in control versus SHHcKI tongues. We found expression of *cell adhesion* genes *(Runx1, Ephb4* and *Fbln2)* were significantly increased in response to SHH (Fig 7B), as were genes common to both GO terms (*Plet1, Pdpn* and *Cxcl10*) although only the increase in *Plet1* was significant (Fig 7C). By contrast, *Tgfb2* was significantly reduced by SHH (Fig 7D). These data support a model where SHH promotes TRC differentiation via regulation of FOXA1 and FOXA2 which in turn regulate genes involved in cell adhesion and cell movement.

## Discussion

Using inducible lineage tracing we show that non-taste lingual epithelium and taste buds have a common embryonic origin from KRT14+ basal cells, and that once placodes have formed, embryonic KRT14+ lingual progenitors do not contribute new cells to taste bud primordia; instead, generation of new TRCs from KRT14+ cells commences at birth and ramps up rapidly in the first 2 postnatal weeks. Additionally, we show that genetic induction of SHH expression represses taste bud development in embryonic tongue epithelium, while at birth SHH promotes taste fate as progenitors are activated. This shift in SHH function does not correlate with the onset of SHH delivery from gustatory neurons, as these neurons express SHH before taste bud primordia are innervated. Using RNAseq to profile SOX2-GFP^high^ taste associated vs SOX2-GFP^low^ expressing lingual epithelial cells, we identified known SHH target transcription factors FOXA2 and FOXA1 enriched in GFP^high^ and GFP^low^ populations, respectively. We find FOXA2 and FOXA1, in addition to known expression of SOX2, are expressed in taste buds at P0 and their expression is altered in response to forced SHH expression. Additionally, we use motif analysis to identify potential targets of SOX2, FOXA1 and/or FOXA2. GO analysis of potential target genes indicate FOXA1;FOXA2 together likely control a suite of genes that function in cell adhesion/migration. Finally, we show expression of these genes is altered by forced SHH expression in vivo.

### Taste placodes and non-taste epithelium share a common KRT14+ progenitor at mid-gestation

In embryos, the surface ectoderm that gives rise to the epidermis is initially a single cell layer of KRT8/KRT18+ cells that by E9.5-10.5 also express KRT14/KRT5, and subsequently lose KRT8/18 expression (Byrne et al., 1994). Extensive lineage tracing studies of KRT14/5+ progenitors has firmly established that the epidermis and its specialized appendages, such as hair follicles and sebaceous glands, derive from this common embryonic population (Pispa and Thesleff, 2003; Thesleff and Mikkola, 2014). Similarly, we show in the developing tongue that KRT8 and KRT14 are broadly co-expressed at mid-gestation and confirm via KRT14 lineage tracing initiated at E12.5 that taste and non-taste epithelia share this common embryonic progenitor. A recent report using inducible Shh lineage tracing initiated at E11.0 identified an early Shh/KRT8 common progenitor population (Kramer et al., 2019), consistent with global epithelial expression of both SHH and KRT8 at this stage (Echelard et al., 1993; Mbiene and Roberts, 2003; Zhu et al., 2014). In contrast to our findings, however, KRT14-Cre lineage tracing presented in the same study resulted in reporter expression only in non-taste lingual epithelium, and not taste placodes or taste bud primordia as late as P1; lineage traced TRCs were only detected at P56 (Kramer et al., 2019)(also see (Bar et al., 2019). It is likely that these different results reflect differences in the efficacy of the *Krt14Cre* alleles employed, as randomly inserted transgenes like the different *Krt14* allele each study employed are notoriously prone to transgenerational silencing (Haruyama et al., 2009). Further, gene promoter-specific transgene silencing can be incomplete, resulting in transgene expression that does not mirror the temporal expression of the endogenous gene product. Here, based on our demonstration of robust KRT14 protein expression at E12.5, we are confident our lineage tracing at this stage with this *Krt14*^*CreER*^ allele (Li et al., 2000) is biologically accurate, and allows us to conclude that taste placodes and non-taste epithelial cells share a common KRT14+ progenitor at mid-gestation.

### KRT14+ taste progenitor competency is activated at birth

Once specified, taste placodes/taste bud primordia do not increase in cell number for the remainder of embryogenesis. This stasis is consistent with numerous reports that taste placodes are mitotically quiescent, while the rest of the lingual epithelium actively proliferates (Farbman and Mbiene, 1991; Liu et al., 2008; Mbiene and Roberts, 2003; Thirumangalathu and Barlow, 2015). Rather we find taste cell number first increases in the first postnatal week, consistent with the demonstrated growth of postnatal taste buds by cell addition (Hendricks et al., 2004; Krimm and Hill, 1998a, b). Importantly we show these new TRCs arise from KRT14+ taste progenitors that are activated at birth, and whose contribution steadily increases with postnatal age. Our data confirm and extend those of a previous report showing postnatal KRT14 lineage labeling of taste bud cells at P9 (Okubo et al., 2009). Although we can pinpoint when progenitors begin to generate new TRCs, it is not known when *bona fide* taste cell replacement begins. In rats, taste buds reach their mature size by P40 (Hendricks et al., 2004), thus input and loss of TRCs must balance by this time. However, we showed previously that over half of SHH+ placode-derived TRCs are lost by P42 in mice (Thirumangalathu et al., 2009). As taste buds continue to enlarge as embryonically derived TRCs are lost, taste bud cell replacement must begin before taste buds reach their mature size. That the rate of production of new TRCs is faster in the first 40 postnatal days than in adults may accommodate the requirement for the growth of postnatal taste buds in the face of steady TRC loss (Hendricks et al., 2004).

### Embryonic SHH+ placodes and postnatal KRT14+ progenitors give rise to all TRC types

In adult mice, KRT14/5+ progenitors give rise to taste cells and non-taste epithelium (Gaillard et al., 2015; Okubo et al., 2009). Taste-fated daughter cells exit the cell cycle, enter taste buds and express *Shh* (Miura et al., 2006). These SHH+ postmitotic taste precursors then differentiate into each of the 3 major TRC types: I – glial-like support cells, II – sweet, bitter or umami detectors, or III – sour sensors (Miura et al., 2014). Here we show that KRT14+ progenitors also produce all 3 TRC types starting at birth. Previously, we found SHH+ placode cells differentiated into type I and II TRCs but did not detect type III cells among *Shh*^*CreER*^ lineage traced taste cells at postnatal stages (Thirumangalathu et al., 2009). Here using a tomato reporter allele, we demonstrate that SHH+ placodes, like adult SHH+ precursor cells are fully competent to produce all TRC types. In addition, we show that embryonic SHH+ placodes and KRT14+ postnatal progenitors produce both type II and III TRCs assessed at early postnatal stages. Thus, we conjecture that both early embryonic and reactivated postnatal KRT14+ progenitors give rise to postmitotic SHH+ cells that in turn differentiate into TRCs; the distinction in embryos is that SHH+ taste primordia remain poised for differentiation for several days until shortly before birth, while adult SHH+ precursors differentiate within 48-72 hour of their terminal division (Miura et al., 2006; Miura et al., 2014).

### SHH has opposing roles in embryonic vs adult lingual epithelium

SHH is a key regulator of taste epithelial development and renewal. In embryos, in vivo and in vitro data indicate that SHH represses taste fate during placode specification and patterning (El Shahawy et al., 2017; Hall et al., 2003; Liu et al., 2004; Mistretta et al., 2003). Here we confirm and expand upon these findings. In addition to more and larger taste papillae and taste buds, we find early genetic deletion of SHH leads to precocious differentiation of TRCs. This is likely due to increased WNT signaling which is normally repressed by SHH to restrain placode size and number (Iwatsuki et al., 2007; Liu et al., 2007). Additionally, excess WNT/ß-catenin signaling during placode specification drives precocious TRC differentiation, even in the absence of Shh (Thirumangalathu and Barlow, 2015), further suggesting Shh deletion at mid gestation leads to upregulated Wnt/ß-catenin signaling which in turn accelerates TRC differentiation. Once placodes are patterned, lingual epithelium is insensitive to Hh pathway inhibition in vitro (Liu et al., 2004), and we confirm this loss of sensitivity in vivo. Following genetic deletion of Shh at E16.5 after placode specification and patterning, taste bud and papilla size do not differ from controls, nor is TRC differentiation accelerated.

In adult tongue, by contrast, SHH promotes TRC differentiation. Genetic deletion or pharmacological inhibition of Hh pathway components in mice inhibits taste cell differentiation (Castillo-Azofeifa et al., 2017; Castillo-Azofeifa et al., 2018; Kumari et al., 2015; Lu et al., 2018), while forced SHH expression induces formation of ectopic taste buds comprising all 3 TRC types (Castillo et al., 2014). Here we show that the pro-taste function of SHH is evident at birth and coincides with the onset of KRT14+ progenitor contribution to taste buds, i.e., forced SHH expression at P0 causes formation of ectopic taste cells. By contrast, forced epithelial SHH expression does not induce TRC differentiation in embryonic lingual epithelium, but as expected, leads instead to smaller and fewer taste primordia, consistent with a recent report where repression of SHH expression in non-taste epithelium has been shown to be essential for taste bud maintenance in late gestation (Bar et al., 2019). Specifically, deletion of PRC-1 chromatin regulator genes in lingual epithelium leads to ectopic SHH expression that is associated with reduced FFP taste buds, and consistent with our study, does not induce ectopic taste buds.

TRC differentiation in adults is maintained by SHH supplied by post-mitotic taste precursor cells within buds and the gustatory innervation (Castillo-Azofeifa et al., 2017; Lu et al., 2018). SHHs role in taste placode patterning is via its epithelial expression as taste bud primordia are only first innervated once placode specification and patterning are complete (Lopez and Krimm, 2006). Nonetheless, we find that subsets of taste fibers already express SHH when lineage tracing is initiated at E12.5 in immature gustatory neurons whose axons have grown into the base of the tongue at this stage but not yet reached their epithelial placode targets (Mbiene and Mistretta, 1997; Scott and Atkinson, 1998). Complete deletion of SHH in gustatory neurons commencing as these neurons are generated has no impact on taste placodes specification or patterning in embryos, nor on taste cell renewal in postnatal mice, indicating that as in adults, epithelial SHH can compensate for loss of a neural supply of SHH to support activation and maintenance of postnatal KRT14+ progenitor function (Castillo-Azofeifa et al., 2017)(but see (Lu et al., 2018).

### Transcriptional profiling implicates FOXA1 and FOXA2 in TRC renewal

The transcription factor SOX2 is required for TRC differentiation in neonates, and high levels of SOX2 have been suggested to be required for progenitor selection of taste over non-taste epithelial fate in adults (Castillo-Azofeifa et al., 2018; Nakayama et al., 2015; Okubo et al., 2006). RNAseq analysis of P0 SOX2-GFP^high^ and GFP^low^ lingual epithelial cells further support this hypothesis as the SOX2-GFP^high^ population is enriched with genes associated with TRC fate. Additionally, SOX2 functions downstream of SHH in adult taste renewal (Castillo et al., 2014; Castillo-Azofeifa et al., 2018), and at birth many SHH pathway components are well expressed in SOX2-GFP^high^ cells. We identified a subset of transcription factors in our dataset linked with the SHH pathway, including FOXA2 and FOXA1, which are known to cooperate with and compensate for one another in development of numerous tissues, including brain and intestine (Ang, 2009; Golson and Kaestner, 2016; Mavromatakis et al., 2011; Metzakopian et al., 2012). As in other developing tissues, the relative expression of these 2 genes in developing lingual epithelium initially overlaps substantially then partially segregates over time suggesting these changes could contribute to changes in the lingual epithelial response to SHH. Consistent with a role in TRC renewal downstream of SHH, we found that ectopic TRCs induced by forced SHH expression also expressed FOXA2, and that epithelial SHH OE promoted FOXA2 and repressed FOXA1 in and adjacent to SHH+ cell patches, thus identifying new candidates that likely coordinately function in TRC differentiation downstream of SHH.

To develop hypotheses as to the role these factors may play downstream of SHH in lingual epithelium, we identified potential gene targets via motif analysis for SOX2, FOXA2 and FOXA1 binding in our list of SOX2-GFP^high^ DEGs. GO analysis revealed that FOXA1/2 may regulate a set of genes associated with cell migration and cell adhesion. All of these genes are enriched in SOX2-GFP^high^ cells (Table 2) and many are associated with cell adhesion changes and movement in both development and disease in a spectrum of tissues (Nieto et al., 2016). Thus, we hypothesize that TRC renewal may involve a partial EMT-MET process as has been proposed in developing and renewing hair follicles (Hong et al., 2015; Jamora et al., 2005; Perez-Moreno et al., 2003). Specifically, as KRT14+ progenitors activate, taste-fated daughter cells must modulate adhesion and motility to allow their exit from the non-taste epithelial adhesive environment to the specialized adhesive environment of taste buds thought to be critical for proper taste bud function (Dando et al., 2015; Michlig et al., 2007). A partial EMT-MET-like behavior has been observed for hair follicle cell renewal from bulge stem cells, where newly generated daughters integrate in distinct cell adhesive environments as they differentiate. The transcription factor OVOL2 is required for this process (Hong et al., 2015; Lee et al., 2017). Intriguingly, *Ovol2* is expressed at low levels in SOX2-GFP^high^ and GFP^low^ cells, while *Ovol1*, which regulates and may have overlapping function with Ovol2 in skin (Teng et al., 2007), is enriched 1.8 fold in P0 SOX2GFP^high^ cells. We also find that *Snai2*, a key EMT gene is well expressed in both GFP^high^ and GFP^low^ populations. While expected in SOX2-GFP^low^, which comprised epithelium and mesenchyme, *Snai2* expression in SOX2-GFP^high^ cells is consistent with the idea that progenitors and taste fated daughter cells activate EMT genes to enter taste buds (see (Wang et al., 2013). Although we must further validate epithelial *Snai2* expression on tissue sections, expression of EMT genes by taste-relevant epithelial cells during TRC renewal may explain how lineage tracing using assumed “mesenchymal” Cre driver alleles results in lineage labeling of subsets of TRCs in adult mice (Boggs et al., 2016; Liu et al., 2012).

## Materials & Methods

### Animals

Male and female mice of mixed backgrounds were maintained and processed at the University of Colorado Anschutz Medical Campus in accordance with approved protocols by the Institutional Animal Care and Use Committee. Mouse lines used include combinations of the following alleles or transgenes: *K14*^*CreER*^ (Li et al., 2000), *Shh*^*CreERT2*^ (Jax 005623), *Phox2b*^*Cre*^ (Jax 016223), *Shh*^*flox*^ (Jax 004293), *R26R*^*Shh-IRES-nVenus*^ (Castillo et al., 2014), *R26R*^*tdTomato*^ (Jax 007914), *R26R*^*YFP*^(Jax 006148), *Sox2*^*GFP*^ (Jax 017592). Embryonic day (E) 0.5 was defined as midday on the day a mating plug was observed. Embryos were recovered on desired days of gestation and staged via the eMouse Atlas Project Theiler criteria (www.emouseatlas.org/emap/ema/home.html).

### Embryonic and neonatal Cre induction

Timed-pregnant dams (intra-peritoneal) or neonatal pups (subcutaneous) received 100 mg/kg of tamoxifen dissolved in corn oil at indicated stages (T-5648, Sigma). *Krt14*^*CreER*^;*R26R*^*tdTomato*^, *Krt14*^*CreER*^; *R26R*^*YFP*^, *Shh*^*CreERT2*^; *R26R*^*tdTomato*^, and *Shh*^*CreERT2/fl*^ mice were induced with a single dose of tamoxifen. *Krt14*^*CreER*^; *Rosa*^*SHHcKI-IRES-nVenus*^ animals and littermate controls received tamoxifen on two consecutive days, beginning at indicated stage.

### Immunostaining

Embryo heads (E12.0-E16.5) or peri- and postnatal tongues (E17.5-P28) were collected in ice cold PBS and fixed for 1hr (light fix) or O/N (hard fix) at 4°C in 2% (vol/vol) paraformaldehyde (PFA) in PBS. Tissue was cryoprotected in 20% (wt/vol) sucrose at 4°C overnight then submerged in OCT (Tissue-Tek, Sakura) and flash frozen on dry ice. 10-week-old Phox2b-ShhcKO and control mice were processed using periodate-lysine-paraformaldehyde (PLP) transcardial perfusion as described in (Gaillard et al., 2015). Serial 12 μm cryosections were collected on SuperFrost Plus slides (Fisher).

For immunostaining, slides were incubated in blocking solution (5% normal goat serum or donkey serum, 1% bovine serum albumin, 0.3% Triton X100 in 0.1M phosphate buffered saline, pH 7.3) for 1 hour at room temperature then incubated with primary antisera in blocking solution overnight at 4°C (2 overnights for transcription factors Sox2, Foxa1, and Foxa2). Sections were rinsed and incubated with secondary antisera in blocking solution for 1 hour at room temperature before nuclear counterstain with Draq5 (Abcam, ab108410) or DAPI (Invitrogen, D3571). Primary antibodies used were: rat anti-Krt8 (1:250; Developmental Studies Hybridoma Bank, TROMA-I/AB_531826); rabbit anti-Krt14 (1:3500; Covance, PRB-155P); mouse IgG2a anti-Ecadherin (1:100; BD Transduction Laboratories, 610181); chicken anti-GFP (1:500; Aves Labs, GFP-1020); rabbit anti-NTPDase2 (1:3000; CHUQ, mN2-36L); rabbit anti-PLCβ2 (1:1000; Santa Cruz, sc-206); rabbit anti-NCAM (1:1000; Millipore, AB5032); goat anti-CAR4 (1:1000; R&D Systems, AF2414); rabbit anti-5HT (1:1000; Immunostar, 20080); rabbit anti-Sox2 (1:100; Cell Signaling, 23064); rabbit anti-Foxa1 (1:100; Abcam, ab170933); rabbit anti-Foxa2 (1:100; Cell Signaling, 3143). Alexa Fluor labeled secondary antibodies were used to visualize staining (1:1000; Thermo Fisher Scientific, A11008, A11010, A21245, A11006, A11081, A21247, A21208, A11039, A21137, A11055, A11056). To facilitate 5-HT detection, 5-Hydroxy-L-tryptophan (Sigma-Aldrich H9772; 80 mg/kg, 6.4 mg/ml in 0.1 M PB, pH 7.4) was injected intraperitonially 1 hour prior to tissue collection. Slides were cover slipped with ProLong Gold mounting medium (Thermo Fisher).

E13.5 and 16.5 tissue collected for whole mount analysis was fixed in 2% PFA for 2 hours at 4°C then stored in PBS at 4°C until processing. Tongues were rinsed in room temperature PBS before undergoing methanol dehydration/rehydration (50% MeOH – 5 min; 70% MeOH – 5 min; 100% MeOH – 15 min; 70% MeOH – 5 min; 50% MeOH – 5 min; 25% MeOH – 5 min). Tissue was washed in 1x PBS with 0.2% Triton-X (PBST, 3x 20-minute washes) then incubated in blocking solution for 2 hours at room temperature (see prior section for composition). Primary antisera incubation in blocking solution occurred for 3 overnights at 4°C with continuous agitation. On day 4, tissue was washed 3x 1hour in PBST then incubated in blocking solution containing secondary antisera and Draq5 nuclear counterstain for an additional 2 overnights at 4°C with continuous agitation. On day 6, tissue was washed 3x 1hour in PBST and mounted on SuperFrost Plus slides (Fisher) with ProLong Gold mounting medium (Thermo Fisher).

### Thymidine analog birthdating

EdU (5-ethynyl-2′-deoxyuridine, Life Technologies) was reconstituted in PBS and administered at 50 μg per gram of body weight to neonatal pups by a single subcutaneous injection. Click-iT Alexa Fluor 546 Kit (Life Technologies) was used to detect incorporated EdU according to the manufacturer’s specifications.

### Image Acquisition and Analysis

Anterior tongues were cut as 12 µm serial sections in 6-slide sets, such that sections on each slide were separate by 72 µm. Approximately 16 sections were cut for each slide across the first ∼1.2 mm of each tongue. After immunostaining, slides were de-identified and taste buds were mapped and quantified across each section/slide using a Zeiss Axioplan II microscope outfitted with a Retiga 4000R camera with Q-Capture Pro-7 software. Images of individual taste buds were acquired using a Leica TCS SP5 II laser scanning microscope with LASAF software and a Leica TCS SP8 laser-scanning confocal microscope with LAS X software. Images of sectioned taste buds were acquired via sequential confocal z-stacks with a z-step interval of 0.75 µm and 11-15 optical sections per z-stack. Immunolabeled cells were tallied by analyzing single optical sections and compressed z-stacks in LASAF, LAS X, and Image J software. All analyses were completed by investigators blinded to condition. KRT8 pixel quantification was conducted using an imstack toolbox developed in MATLAB as previously described (Castillo-Azofeifa et al., 2017). For taste bud innervation analysis, taste buds were classified as “innervated” when at least one SHH-tomato+ nerve fiber was present within the FFP papillae core and it reached an apically situated tomato+/KRT8+ taste bud. When quantifying neurites innervating a given taste bud, all tomato+ neurites within the papilla core were counted.

For whole mount E13.5 and E16.5 taste placode analysis, anterior tongues were imaged on a Leica TCS SP5 II laser scanning microscope with LASAF software. Taste placode maps were created from compressed sequential z-stacks consisting of 25 optical sections with a 0.5 µm z-step interval taken at 10x. Each KRT8+ taste placode within the first 1mm of the dorsal surface of the anterior tongue was individually numbered, and 10 taste placodes/tongue were chosen at random (random.org) for quantification. For placode cell number and volume analysis, individual E13.5 and E16.5 taste placodes were imaged from their dorsal to ventral extent at 63x with a 1.5x digital zoom and a micro-sampling z-step interval of 0.25 µm. Placode z-stacks were analyzed in Imaris 8.4 (Oxford Instruments), using the ImarisCell tool. Briefly, a cube-shaped ROI was set in the x, y, and z planes using the KRT8 channel expression as a guide. E-Cadherin expression was utilized by the software to delineate individual cell membranes within the ROI. KRT8 co-expression was then used to define intra-placodal versus extra-placodal cells. Each rendering was de-identified and manually reviewed to assure that cell boundaries were accurately delineated.

### RNA Extraction and qPCR

Freshly harvested tongues were rinsed in chilled in 1x dPBS. To isolate anterior tongue epithelium, enzyme peeling solution (1:1 mix of 2.5% pancreatin in L15 medium and 0.25% Trypsin EDTA) was injected beneath the dorsal and ventral lingual epithelia and tongues incubated at room temperature for 45-60 minutes prior to manual dissection. Postnatal enzyme peeling solution contained 5 mg/mL Dispase II + 3 mg/mL Collagenase II in 1X dPBS and tongues were incubated for 31 minutes at room temperature prior to manual dissection. Peeled tissue was snap frozen on dry ice and stored at −80°C.

Total RNA was extracted with a RNeasy Plus Micro kit (Qiagen, 74034) according to the manufacturer’s instructions, and RNA quantity measured by Nanodrop (ThermoFisher Scientific). mRNA was reverse transcribed using the iScript cDNA synthesis kit (Bio-Rad). SYBR Green-based qPCR was performed by using Power SYBR Green PCR Master Mix reagent (Applied Biosystems) and gene-specific primers (*see table*). qPCR reactions were carried out in triplicate using a StepOne Plus Real-Time PCR System (Applied Biosystems, Life Technologies). Relative gene expression was analyzed using the ΔΔCT method (Livak and Schmittgen, 2001). The ribosomal gene *Rpl19* was used as an endogenous reference gene.

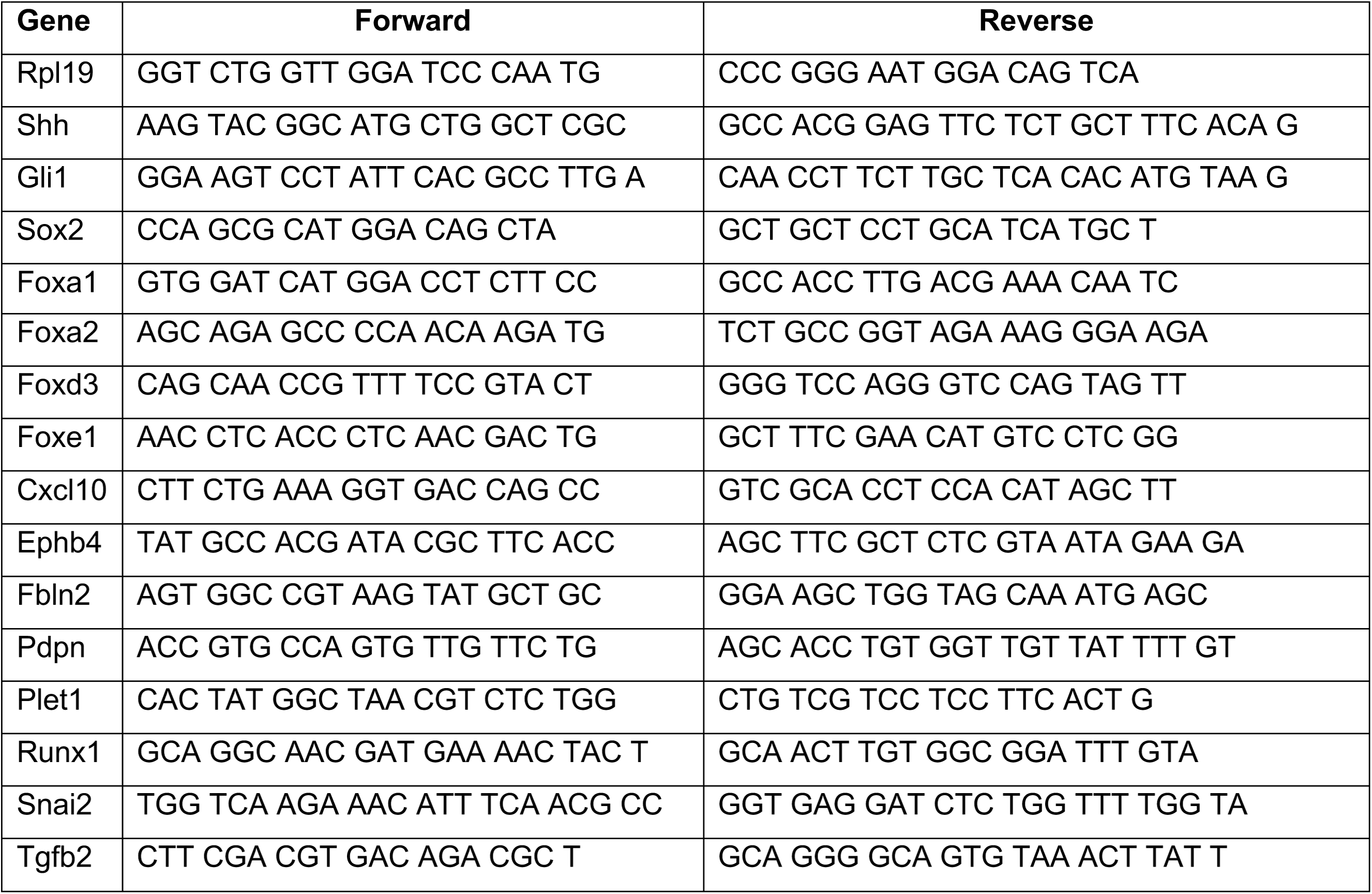

### Fluorescence-Activated Cell Sorting (FACS) and RNA Extraction for RNA-Seq

Lingual epithelium was isolated from SOX2-GFP P0 mice using embryonic enzyme peeling solution and manual dissection as described above (45-60 min room temperature incubation in 1:1 mix of 2.5% pancreatin and 0.25% Trypsin EDTA). Peeled tissue pieces were chopped with a sharp blade then further digested in enzyme solution of 1 mg/mL Collagenase II and 1 mg/mL Elastase in 1X dPBS for 1hr at 37°C. Digested tissue was manually triturated into single cells using a sterile glass pipette then pelleted in DMEM/10% FBS to inhibit further enzyme digestion. Pellets were resuspended in FACS Buffer (1mM EDTA, 25 mM HEPES pH 7.0, 1% FBS, and 1x Ca^2+^/Mg^2+^ free dPBS) and passed through a 30 μM nylon mesh filter to isolate single cells from remaining aggregates. FACS-based purification of SOX2-GFP cells was carried out on a MoFlo XDP70 (Gates Center Flow Cytometry Core, University of Colorado Anschutz Medical Campus) according to the green fluorescent protein signal (excitation 488nm; emission 530 nm). Red fluorescent protein channel (582 nm) was used to gate out autofluorescent cells, and DAPI (450 nm) was used to gate out dead cells. Total RNA was extracted from FACS isolated cell pools (GFP^high^, GFP^low^, GFP^neg^) using the Arcturus PicoPure RNA Isolation Kit (Thermo Fisher #KIT0204) according to manufacturer’s instructions.

### RNA-Seq Analysis

Illumina HiSEQ libraries were prepared and sequenced by the Genomics and Microarray Core Facility at the University of Colorado Anschutz Medical Campus (Illumina HighSEQ4000, HT Mode, 1×50). Base calling and quality scoring were performed using Illumina Real-Time Analysis (RTA) v2.7.3 software. Files were demultiplexed and converted from BCL to FASTQ format using bcl2fastq2 v2.16.0.10 conversion software. Single-end reads were trimmed using Trimmomatic v0.36 and subsequently mapped to the mm10 mouse genome using GMAP/GSNAP v20141217. FPKM values were then obtained using Cufflinks v2.2.1. Analysis was conducted using R 3.5.3. Non-protein coding genes and genes with no expression across all samples were removed prior to analysis. Genes with an absolute fold change greater than or equal to 1.5 and FPKM values greater than or equal to 5 in each sample were designated as differentially expressed (DEG).

### Gene Ontology (GO) analysis

Gene Ontology (GO) analysis of differential expressed genes (DEGs) was performed in g:Profiler using user default settings (Tables 4 and 5) and GO enrichment analysis powered by PANTHER (geneontology.org) (Tables 8 and 9).

### Motif enrichment analysis

Regions of interest (ROIs) that included 1 KB upstream and 200 bp downstream of the transcription start site were extracted for the 1032 differentially expressed GFP^high^ genes using the ‘biomaRt’ package for R. For genes with multiple isoforms, the APPRIS principle isoform was selected. In the few cases where multiple principle isoforms were annotated with APPRIS, the isoform with the transcription start site most upstream of the gene start site was used, resulting in one ROI for each differentially expressed gene. ROIs were converted to BED format and then FASTA format using a custom PERL script and the Ensembl GRCm38 mouse genome (release 96).

FASTA files of ROIs were used as the input for AME (MEME-Suite, v5.0.5) and were tested against the HOCOMOCO mouse V11 motif database using default parameters but for the ‘--control -- shuffle--’ flag. AME outputs were parsed with a custom R script to compare motif enrichment and gene expression data. Data are presented as the ratio of motif hits in the primary sequence divided by motif hits in the background (shuffled primary sequences).

### Statistical Analyses

Normally distributed data were analyzed in Graphpad Prism 8 software using parametric two-tailed Student’s *t*-tests. Significance was taken as p < 0.05 with a confidence interval of 95%. Data are presented as mean ± standard deviation (SD) unless otherwise indicated.

## Acknowledgements

The authors would like to thank the staff of the Gates Center for Regenerative Medicine Flow Cytometry Core and the staff of the University of Colorado Cancer Center Genomics and Microarray Shared Resource, as well as Kelly Zaccone and Ernesto Salcedo for technical support. Thank you to Fred DeSauvage, Genentech for providing the initial ShhcKI-YFP mice. Thank you also to our colleagues in the Rocky Mountain Taste and Smell Center as well as the Department of Craniofacial Biology at the University of Colorado Anschutz Medical Campus for constructive conversations and feedback throughout the duration of the study. This work was support by grants from the National Institutes of Health/National Institute for Deafness and Other Communication Disorders to EJG (F32 DC015958) and to LAB (R01DC012383), and from the National Institutes of Health/Cancer Center Support Grant to the Gates Center of Regenerative Medicine (P30CA046934).

## Supplementary Figure Legends

**Figure S1. Taste receptor cell differentiation begins before birth.** (**A, B**) NTPDase2+ (green) type I TRCs are detected in taste bud primordia at E17.5 and taste buds at P2. (**C**) The proportion of taste buds housing type I cells increases around birth, such that ∼100% contain NTPDase2+ cells at P4. (**D, E**) PLCß2+ (green) type II cells are detected in taste bud primordia and immature buds at E17.5 and P4, respectively. (**F**) The proportion of taste buds housing type II cells increases to ∼100% by P4. (**G, H**) Small numbers of NCAM+ (green) type III cells (arrowheads), in addition to NCAM+ gustatory neurites (yellow arrows), are detected in taste bud primordia and immature taste buds at E17.5 and P2, respectively. (**I**) Type III cells are detected rarely before birth but found in ∼50% of buds by P2. (A, B, G, H) single 0.75 um optical sections; (D, E) compressed Z-stacks (11 x 0.75 µm optical sections). Draq5 nuclear counterstain (magenta). Dashed lines delimit FFP core and encircle taste primordia (A, D, G) or buds (B, E, H). N=3-4 mice per time point, 65-120 taste buds per animal.

**Figure S2. The timing of genetic deletion of Shh determines its impact on taste development in vivo. (A)** *Shh* and *Gli1* are reduced in tongues of *Shh*^*CreER/fl*^ embryos at E18.5, induced at E13.5. **(B)** Shh deletion at E12.5 results in more KRT8+ taste buds at E18.5, but Shh deletion at E16.5 does not alter taste bud number. (N= 3-5 embryos per timepoint per genotype; Students *t*-test **p < 0.01) **(C, D)** Genetic deletion of Shh at E12.5 increases KRT8+ (green) taste bud profile diameter (yellow dash with arrows) and FFP diameter (white dash). Dotted line indicates basement membrane. Scale bar: 10 μm. (**E, F**) Taste bud and papilla diameter increases significantly in *Shh*^*CreER/fl*^ induced at E12.5 and assayed at E18.5. (N=3-4 embryos per genotype, 20 taste buds per animal, Students t-test, *** p < 0.001, ** p< 0.01). (**G-I**) The proportion of E18.5 taste buds housing type I (NTPdase2+), II (PLCß2+) and III (NCAM+) TRCs is significantly increased by Shh deletion at E12.5. (N=5-8 animals per genotype, Student *t-*test, * p < 0.05) (**J, K**) Shh deletion at E16.5 does not change taste bud number or size assayed at E20. **(L-N)** Shh deletion at E16.5 does not lead to precocious differentiation of type I (NTPdase2+), II (PLCß2+) and III (NCAM+) TRCs at E20. Black bars: mean ± SD for all plots.

**Figure S3. Forced expression of SHH in embryonic lingual epithelium represses development of endogenous taste buds but does not induce ectopic taste buds. (A-D)** Tamoxifen induction of K14-ShhcKI-YFP at E12.5 leads to mosaic YFP expression at P14 (**A, B:** green), but ectopic KRT8+ taste buds do not form in these patches (white **C, D, E**). Endogenous FFP KRT8+ taste buds are detected (**C**: asterisk) but are reduced in number in mutants compared to controls (**E**). (**A, B, F, G**) SOX2 expression (magenta **A, B**; white **F, G**) appears upregulated (arrowheads **F, G**) in the larger ShhcKI-YFP+ patches in non-taste epithelium (white **H, I**), consistent both with the upregulation of SOX2 by ectopic SHH in adult tongues (Castillo et al., 2014) and the report that overexpression of SOX2 in embryonic lingual epithelium in insufficient to drive taste bud differentiation (Okubo et al., 2006). Scale bar: 10 μm.

**Figure S4. Summary of SOX2-GFP^high^ vs SOX2-GFP^low^ differentially expressed genes. (A)** A scatter plot of the Log10 distribution of DEGs enriched in each subpopulation. Fold enrichment of the top 50 genes enriched in (**B**) SOX2-GFP^high^ and (**C**) SOX2-GFP^low^ P0 lingual epithelial cells.

**Figure S5. Changes in expression of SOX2, FOXA2, and FOXA1 in developing FFP and taste buds may underlie changes in the lingual epithelial response to HH signaling. (A-C)** <u>E13.5.</u> **(A)** SOX2 (green) is highly expressed in specified taste placodes (KRT8, magenta, asterisk) and more dimly expressed in adjacent keratinocytes (white arrowheads). FOXA2 (green) (**B**) and FOXA1 (green) (**C**) are expressed at comparable levels by taste placodes (asterisk) and non-taste epithelium (white arrowheads). (**D-F**) <u>E16.5.</u> (**D**) SOX2 is more highly expressed by KRT8+ taste bud primordia (magenta) and adjacent perigemmal cells (yellow arrows), with lower expression in FFP and non-taste epithelium (white arrows and arrowheads, respectively). (**E**) FOXA2 persists in KRT8+ taste primordia (magenta) but not perigemmal or apical FFP epithelial cells (yellow arrowheads); low expression is detected in basal FFP keratinocytes (white arrows) and non-taste epithelium (white arrowheads). (**F**) FOXA1 is lowly expressed by subsets of cells in KRT8+ taste primordia (magenta), is absent from perigemmal and apical FFP epithelial cells (yellow arrowheads) but is expressed in FFP basally (white arrows), developing filiform papillae (fi) and interpapillary epithelial cells (ip). (**G-I**) <u>P0.</u> (**G**) SOX2 remains strongly expressed in KRT8+ taste buds and perigemmal cells (yellow arrows), is detected at lower levels in FFP basal keratinocytes (white arrows) and expressed dimly by non-taste basal cells (white arrowheads). (**H**) FOXA2 expression is restricted to KRT8+ taste buds (magenta) and FFP basal cells at the base of the papilla (white arrows), and absent from perigemmal and apical FFP cells (yellow arrowheads) and non-taste epithelium. (**I**) FOXA1 is evident KRT8+ taste buds, cells at the base of FFP (white arrows), and absent from apical FFP epithelium and perigemmal cells (yellow arrowheads). Epithelial surface indicated by solid line; dashed lines delimit basement membrane. pc: papilla core. Scale bar: 10 μm.

