## Supplementary figures and images for "Onset of taste bud cell renewal starts at birth and coincides with a shift in SHH function"

### Suppl Fig 1

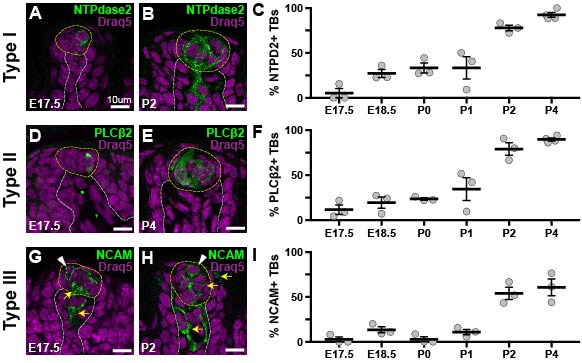

### Suppl Fig 2

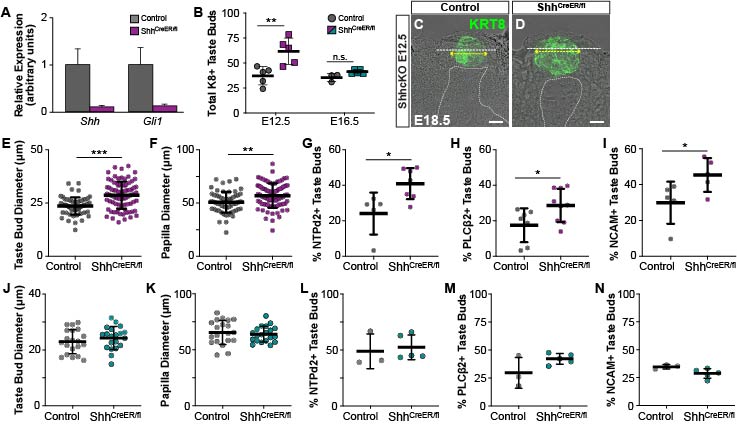

### Suppl Fig 3

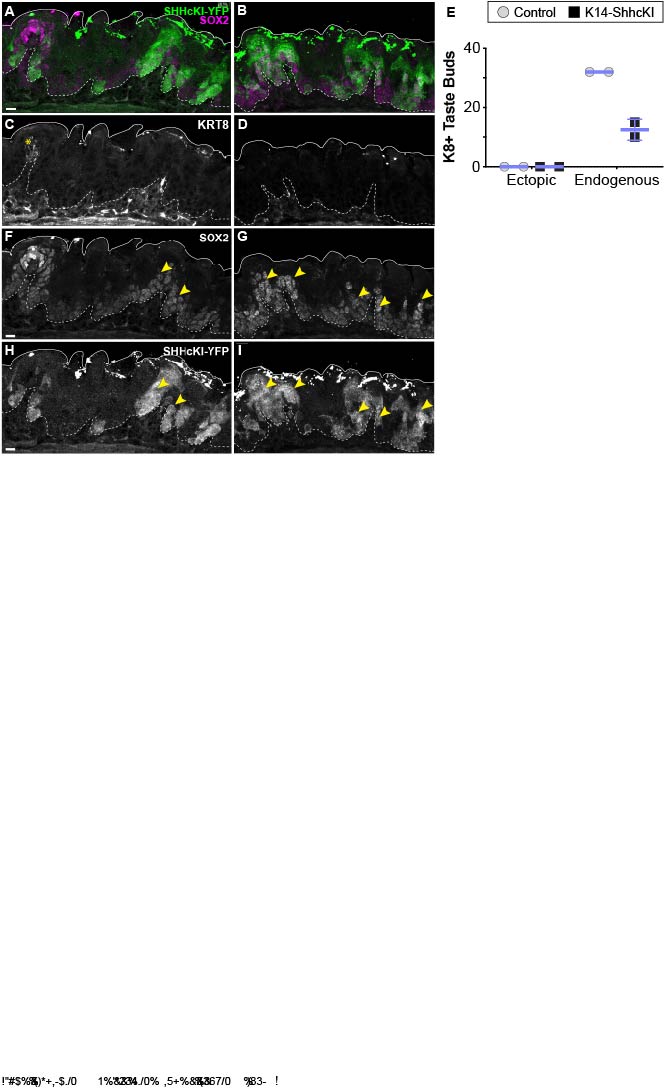

### Suppl Fig 4

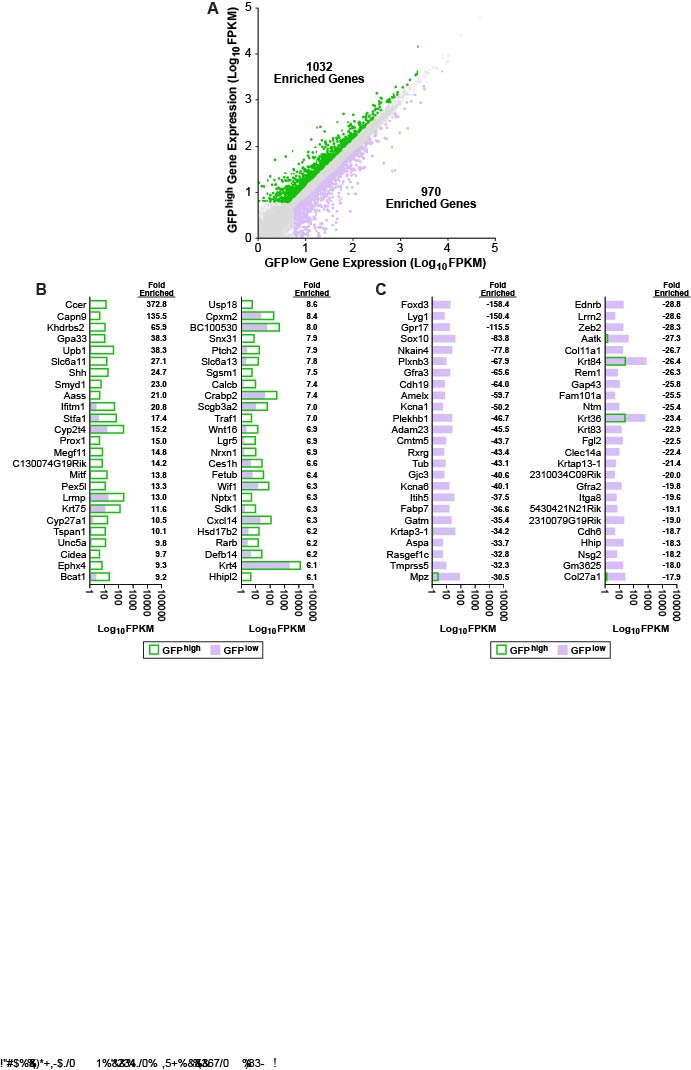

### Suppl Fig 5

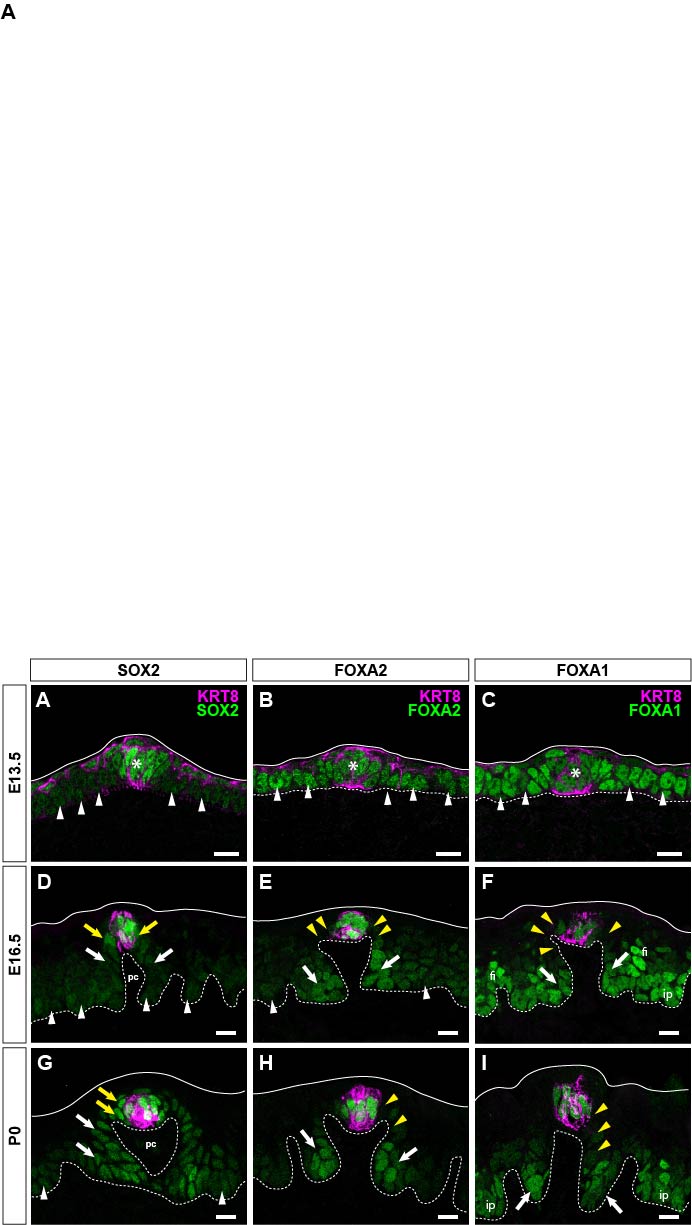
